# Capsular polysaccharide can sensitize bacteria to non-antibiotic drugs

**DOI:** 10.64898/2026.07.28.741313

**Authors:** Matthew O. Gill, Jane A. Cook, Kevin X. Jiang, Achuthan Ambat, Handuo Shi, Aravind Natarajan, Rachael B. Chanin, Zachary Benmamoun, Weifeng Lin, Danica T. Schmidtke, Eric T. Kool, Gavin Sherlock, Lynette Cegelski, Kerwyn Casey Huang, Ami S. Bhatt

## Abstract

Medications administered over long durations, such as phenothiazine antipsychotics, accumulate in the gut at concentrations that affect microbial growth. However, the bacterial features influencing sensitivity to these non-antibiotics remain poorly understood. Bacterial capsular polysaccharides (CPSs) typically confer protection against environmental stressors, including chemical, viral, and immunological pressures within the gut. But their roles under non-antibiotic drug pressure are unknown. Here, we show that the K5 CPS of *Escherichia coli* Nissle 1917 (*EcN*) sensitizes it to thioridazine (TDZ) and related antipsychotics. Among a panel of *E. coli* strains grown in minimal medium, *EcN* exhibited the highest sensitivity to TDZ. Experimentally evolving *EcN* under gut-relevant TDZ concentrations selected for resistant populations with convergent variants affecting the CPS locus, and TDZ-resistant clones correspondingly had lower CPS expression than their drug-susceptible counterparts. Genetic, transcriptomic, and phenotypic analyses confirmed that the K5 CPS enhances, rather than mitigates, TDZ sensitivity. These findings demonstrate that canonically protective surface structures can become vulnerabilities under non-antibiotic pharmaceutical pressure. Human medications may therefore inadvertently shape the expression and evolution of bacterial surface structures in the gastrointestinal tract, challenging presumptions of CPS-mediated environmental protection.

## Introduction

Medications are the most potent modifiers of the human gut microbiota in industrialized populations, exerting effects that exceed those of diet and lifestyle.^1,2^ Human protein-targeted drugs (hereafter referred to as non-antibiotics) disrupt microbial communities through multiple mechanisms: they alter host gastrointestinal (GI) physiology^3,4^, accumulate as substrates for microbial metabolism within the lumen^5,6^, and, in some cases, directly inhibit bacterial growth.^7–11^ Although several non-antibiotics have activity against gut commensals, the bacterial features that govern susceptibility to these compounds remain poorly understood.

This mechanistic knowledge gap is exemplified by antipsychotics, a class of drugs that exhibit broad antibacterial activity despite targeting human neurotransmitter receptors, which lack bacterial homologues.^7,12,13^ Because these medications are typically taken chronically over years to decades and are excreted predominantly via biliary-fecal routes rather than renally^14–16^, they reach sustained and elevated concentrations in the colon. Consistent with this pharmacokinetic profile, several clinical studies have reported associations between antipsychotic use and profound shifts in microbiome diversity and composition.^17–20^ Thus, in line with their direct antibacterial activity, these non-antibiotic drugs might drive bacterial adaptation under chronic exposure.

To dissect these responses mechanistically, we focused on phenothiazines (e.g. TDZ, chlorpromazine), a highly active subclass of antipsychotics that are enriched for antibacterial activity against gut commensals^7,12^, and robustly remodel *in vitro* gut communities.^8,21,22^ Their antimicrobial effects were recognized decades ago^23^, and despite known side effects like QTc prolongation^24^, they remain under investigation for potential clinical use^25–28^, and have been used on a compassionate-care basis for the treatment of multi-drug resistant tuberculosis.^29,30^ While their bactericidal mechanisms are relatively well defined, encompassing membrane permeabilization, proton motive force collapse, and inhibition of efflux pumps and ATP synthesis^26,31^, their variable efficacy - particularly across Gram-negative taxa - remains unexplained.^7,12,32,33^ This paradox suggests that strain-specific determinants, such as surface structures, may govern susceptibility beyond drug potency alone.

To identify phenothiazine susceptibility determinants and map the evolutionary consequences of phenothiazine exposure, we leveraged *Escherichia coli*, a genetically tractable Gram-negative enterobacterium with extensive phenotypic heterogeneity.^34–36^ A screen of several *E. coli* strains against TDZ, a representative phenothiazine antipsychotic, revealed growth of *E. coli* Nissle 1917 (*EcN*) to be the most robustly inhibited, with sensitivity strongly modulated by growth conditions. Guided by this vulnerability, we experimentally evolved *EcN* under physiologically relevant TDZ concentrations^14,37^ to identify candidate adaptive mutations in TDZ-resistant populations and clones. Our analyses revealed that the K5 capsular polysaccharide (CPS), classically viewed as a protective barrier^38–41^, paradoxically confers maximal drug susceptibility. Further testing confirmed that CPS-mediated sensitivity extends to other phenothiazine antipsychotics, highlighting the capsule as an unexpected vulnerability factor under non-antibiotic selective forces. These findings place non-antibiotic drugs alongside phages and host immunity as potent selective forces shaping the interface of gut-associated bacteria with their environment.^42^

## Results

### EcN exhibits strong susceptibility to TDZ despite robust efflux capacity

Given the variable susceptibility of Gram-negative bacteria to phenothiazine antipsychotics,^7,12,32,33^ we assembled a panel of four diverse *E. coli* strains (**Fig. 1a**). The panel comprised a laboratory strain (K-12 BW25113), the colon-colonizing probiotic (Nissle 1917, *EcN*)^43,44^, a bovine diarrheal strain (ATCC 31616), and a human neonatal meningitis-associated strain (ATCC 700973).

**Figure 1.**
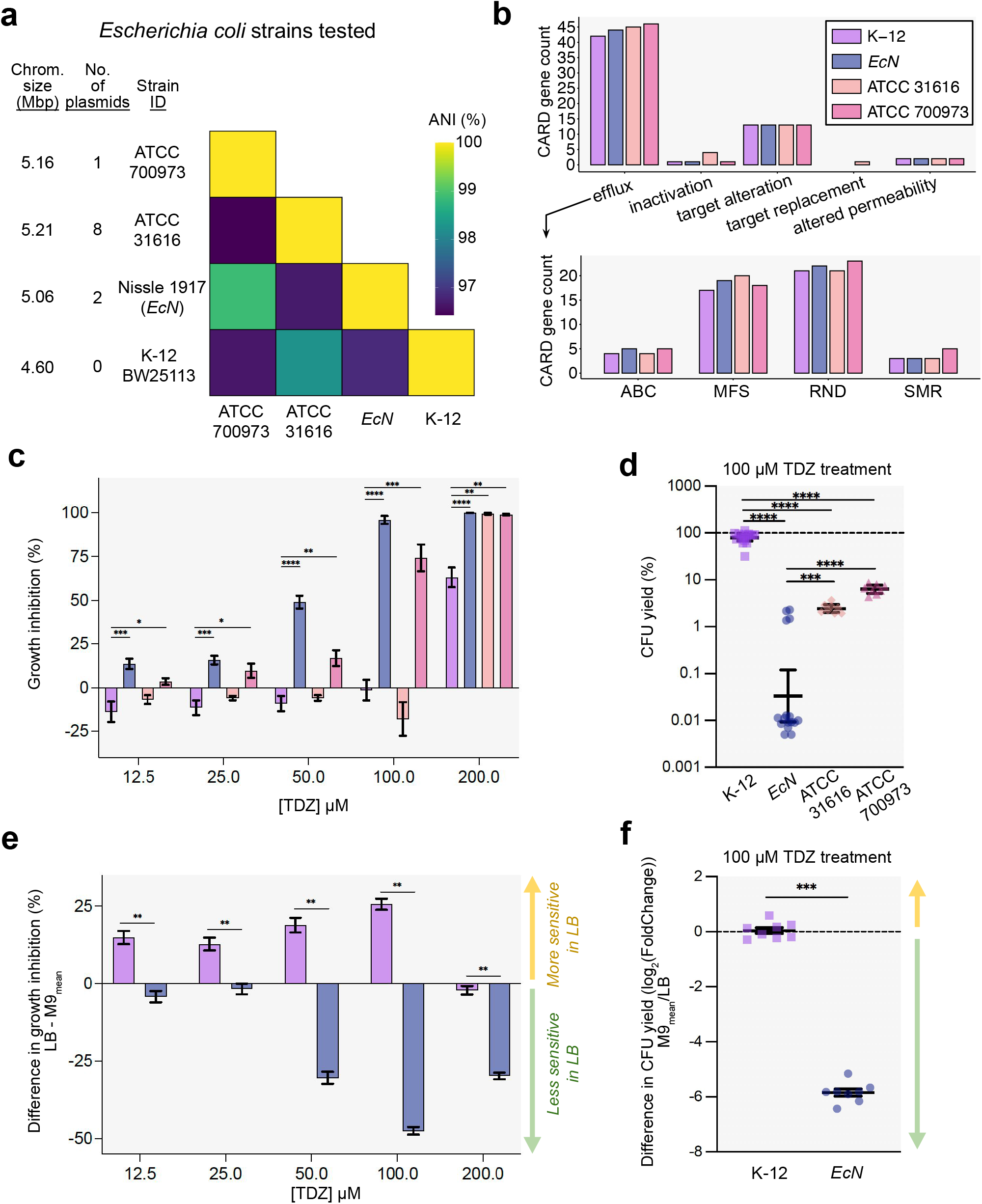
*E. coli* Nissle 1917 exhibits strain-specific sensitivity to TDZ that is minimized in rich media. (**a**) Genome size, plasmid number, and average nucleotide identity (ANI %) across four *E. coli* strains screened for drug sensitivity. (**b**) Antibiotic resistance gene counts. Top panel: functional categories. Bottom panel: efflux pump superfamilies (ATP-Binding Cassette (ABC), Major Facilitator Superfamily (MFS), Resistance-Nodulation-Division (RND), Small Multidrug Resistance (SMR)). (**c**) Dose-dependent growth inhibition (%) of *E. coli* strains by TDZ in M9-glucose, assessed by endpoint OD_600_ assays. Data are mean ± SEM from a minimum of two biological (four for K-12 and *EcN*) and two technical replicates. K-12 was compared individually against each other strain. (**d**) CFU yield (%), relative to vehicle treatment, after five hours of 100 µM TDZ treatment initiated at exponential phase. Error bars are the 95% CI centered on the geometric mean. Data are from a minimum of two biological (four for K-12 and *EcN*) and four technical replicates. K-12 and *EcN* were compared individually against each other strain. (**e**) Difference in growth inhibition (%) in LB broth compared to M9-glucose. Growth inhibition (%) in LB was first calculated from two biological and three technical replicate cultures, as was previously done in M9-glucose (panel C). Negative values reflect less inhibition in LB compared to M9. (**f**) Log_2_(Fold-Change) in CFU yield (%) for two biological and four technical replicates in LB compared to the mean CFU yield (%) in M9-glucose. Error bars are SEM, centered on the mean. Negative values reflect higher yield in LB compared to M9-glucose. Mann-Whitney U was used for all statistical comparisons: * = p < 0.05, ** = p < 0.01, *** = p < 0.001, **** = p < 0.0001. Absence of a reported significance value indicates a non-significant difference.

Previous work established that, in *E. coli* K-12, the outer membrane channel *tolC* is required for maximal resistance to 40 µM TDZ.^7^ We confirmed and extended these findings using a vehicle-normalized OD_600_ endpoint growth assay: the Δ*tolC* K-12 strain exhibited significantly enhanced sensitivity compared to wild type across a broad range of TDZ concentrations (12.5–200 µM) in both rich and minimal media (**Fig. S1**). Given the central role of efflux pumps in TDZ resistance, we next hypothesized that differences in efflux pump copy number might predict differential sensitivity to TDZ.^46^ We interrogated publicly available and ATCC reference genomes using the *Resistance Gene Identifier* from the *Comprehensive Antibiotic Resistance Database* (CARD)^47^, to quantify antibiotic resistance determinants, focusing on efflux pumps. This analysis revealed minor variations in efflux gene counts across the panel with K-12 encoding the fewest total efflux pump genes, differing from other strains by four copies or fewer (**Fig. 1b**). Consequently, we predicted that TDZ sensitivities would be broadly similar across isolates.

To test whether these strains were differentially sensitive to TDZ, we next screened the four isolates in M9-glucose minimal medium, as nutrient limitation can amplify vulnerability factors that are silent under rich growth conditions.^48^ Contrary to our prediction, strain-specific susceptibility to TDZ varied greatly. Under the nutrient-limited conditions, both *EcN* and ATCC 700973 were more sensitive to TDZ than K-12 at all TDZ concentrations tested (**Fig. 1c**). At 50 µM and 100 µM – approximating conservative estimates of small intestine (∼40 µM) and colon (∼100 µM) TDZ concentrations (see **Methods**)^14,37^ – *EcN* was the most vulnerable strain, displaying 49% and 96% growth inhibition, respectively (**Fig. 1c**). We orthogonally evaluated these findings with a complementary colony-forming unit (CFU) viability assay. When exposed to 100 µM TDZ for five hours starting at mid-exponential phase (OD_600_∼0.3), *EcN* exhibited drastically reduced viability (median 0.011% CFU yield) compared to all other strains (**Fig. 1d**). ATCC 31616 showed lower CFU yield than K-12 despite appearing largely uninhibited in the OD_600_ assay (**Fig. 1c**). Collectively, these data confirm that, while efflux pumps are essential for *E. coli* survival at physiologically relevant TDZ concentrations (**Fig. S1**), efflux pump encoding alone cannot predict strain-specific susceptibility to TDZ.

### EcN susceptibility to TDZ is potentiated in minimal medium

The pronounced sensitivity of *EcN* (**Fig. 1c-d**) suggests the presence of a strain-specific determinant that enhances TDZ vulnerability. *EcN* has over 1,500 unique coding sequences (CDS) compared to K-12 MG1655^49^, with significant enrichment of surface-associated and virulence-related proteins – including siderophores, adhesins, type VI secretion systems, toxin-antitoxin modules, and the K5 CPS – which are thought to promote the intestinal engraftment and probiotic properties of *EcN*.^38,49–52^ Metabolism-linked genes (>300 CDSs) are also enriched in *EcN*.^49^ Given that both gene categories can be regulated by nutrient availability^53–55^, we hypothesized that *EcN*’s strain-specific sensitivity would be affected by growth in nutrient-replete conditions. Consistent with this hypothesis, *EcN* was markedly less susceptible to TDZ in LB broth at concentrations ≥50 µM (**Fig. 1e**). By contrast, K-12 was more sensitive when cultured in LB compared to M9-glucose across nearly all tested concentrations (**Fig. 1e**). CFU viability measurements of mid-exponential phase cultures confirmed that LB specifically attenuates *EcN*’s susceptibility to 100 µM TDZ, whereas K-12’s susceptibility remained unaffected (**Fig. 1f**). Together, these data demonstrate that *EcN*’s TDZ sensitivity is strain-specific and strongly medium dependent.

### Colon-approximate TDZ exposure drives rapid adaptation and stable resistance

To investigate the genetic basis of strain-specific and medium-dependent TDZ sensitivity in *EcN*, we turned to experimental evolution. Given that we found *EcN* to be severely inhibited by TDZ at colon-approximate concentrations (**Fig. 1c-d**), we hypothesized that 100 µM-exposed populations might accrue resistance to TDZ through either efflux pump upregulation^56–59^, mutations in *EcN*-specific vulnerability factors, or a combination of the two. We experimentally evolved replicate *EcN* populations in M9-glucose under three conditions: vehicle-only (“pDMSO”), small intestine-approximate (40 µM; “p40”), or colon-approximate TDZ concentrations (100 µM; “p100”) (**Fig. 2a**). We assessed population-level adaptation across multiple passages using standard growth curve-derived fitness metrics (**Fig. S2a**)^60,61^, defining adaptation by the ability to grow in 100 µM TDZ and outright resistance by sustained growth in 200 µM TDZ (based on initial screening; **Fig. 1c-d**).

**Figure 2.**
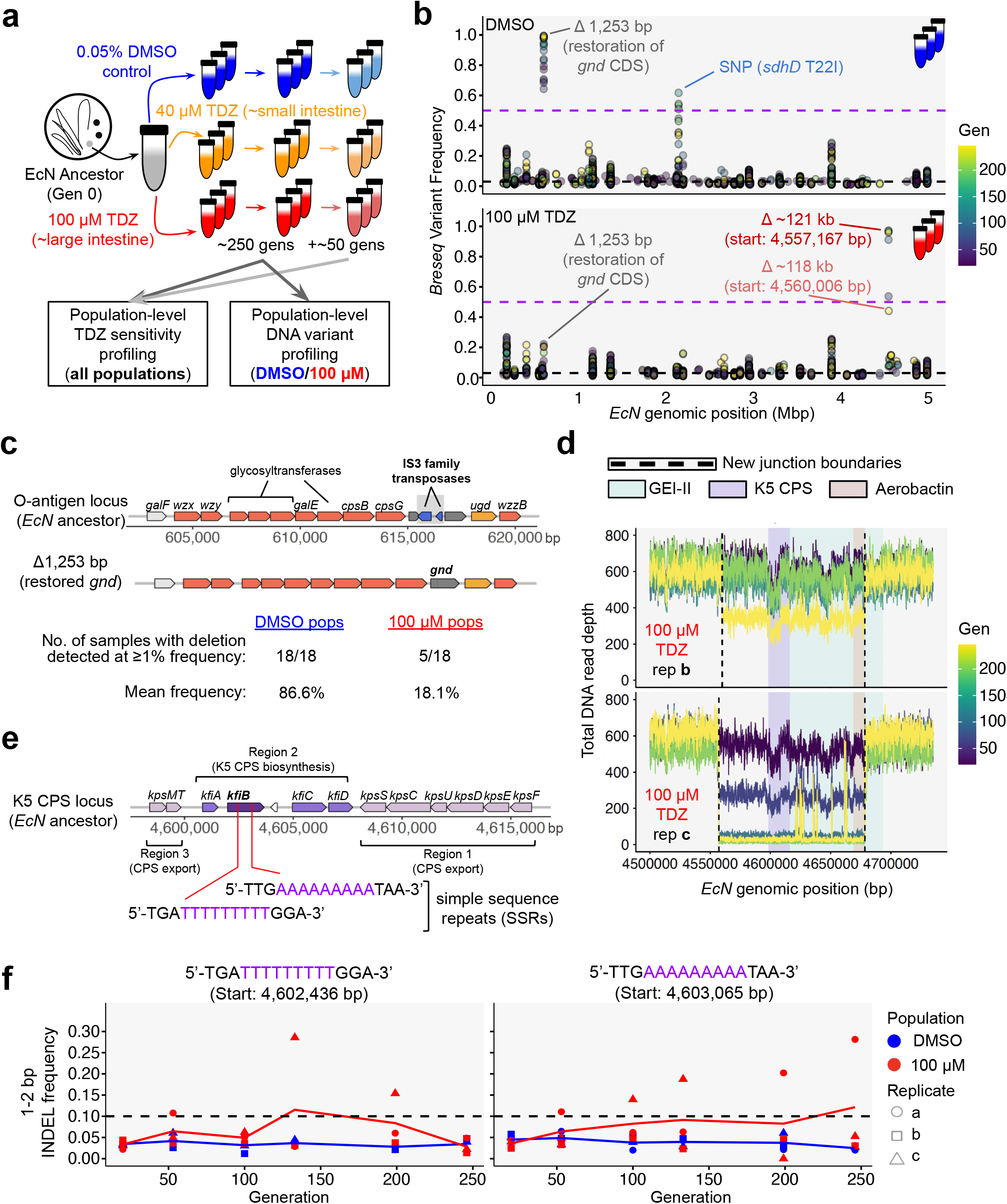
TDZ-adapted populations have structural variations and reversible mutations affecting surface polysaccharide loci. (**a**) Overview of the experimental evolution. Gens = generations. (**b**) Frequencies of *breseq*-detected variants in the DMSO (top panel) and 100 µM TDZ (bottom panel) replicate populations across six sequenced generations (20, 53, 100, 133, 199, 246). Only variants reaching ≥3% (black dashed line) in at least one population at any timepoint were included in the analysis; variants below the minimum detected frequency (1%) were omitted from the plot. Variants breaching 50% frequency (purple dashed line) uniquely in either pDMSO or p100 were flagged as variants of interest. (**c**) The O-antigen locus in our ancestral, wild-type *EcN* genome, which harbors an IS element (ISEc52) that truncates *gnd*. Genes in dark orange are linked to O-antigen/LPS biosynthesis in *EcN*. The number of replicates across all timepoints that harbored the ISEc52 “deletion” variant at ≥1%, and their mean frequency in pDMSO and p100 populations separately, are below. (**d**) Total read depth across the unique ∼120 kb deletion loci detected in p100 replicate *b* (top) and replicate *c* (bottom). A previously described genomic island (GEI-II), the K5 CPS, and aerobactin loci are highlighted. (**e**) The K5 CPS locus in our ancestral, wild-type *EcN* genome. The *kfiB* gene sequence contains two SSRs subject to early stop-inducing INDELs uniquely in the p100 populations. (**f**) Frequencies of 1-2 bp INDELs in the two *kfiB* SSRs that reached ≥10% frequency (black dashed line) in at least one replicate at any timepoint.

Exposure to 40 µM TDZ failed to select for drug-specific population-wide adaptation: across all passages, p40 cultures remained indistinguishable from pDMSO in vehicle-normalized AUC, carrying capacity, and growth rate (**Fig. S2b-d**). Occasional low-level growth in 200 µM TDZ at later passages (generations 199 and 246) was sporadic and observed in both p40 and pDMSO replicates, indicating it was not drug-selected (**Fig. S2b-d**). By contrast, 100 µM TDZ rapidly drove full resistance. All p100 populations grew robustly in 100 µM TDZ by generation 100, with vehicle-normalized fitness metrics in 200 µM TDZ approaching 1.0 by generation 246 (**Fig. S2b-d**). This resistance persisted through ∼53 additional generations of drug-free passaging, demonstrating that colon-approximate TDZ concentrations select for stable, heritable resistance in *EcN* populations.

### TDZ-resistant populations harbor convergent variants in surface polysaccharide loci

To identify adaptative mutations, we sequenced pDMSO and p100 population replicates at six timepoints (20, 53, 100, 133, 199, and 246 generations) and called variants using *breseq* (**Methods**)^62^. We focused on mutations that were uniquely detected with ≥50% frequency in only one treatment group, identifying four key variants – three of which converged on surface polysaccharide-linked loci: the O-antigen-associated *gnd* locus and the K5 CPS locus (**Fig. 2b**).

The first variant is a 1,253 bp deletion within the O-antigen cluster that restores an intact *gnd* open reading frame. Despite reaching a maximum frequency of 22.1% in one p100 population replicate, it was only detected (i.e., reaching a minimum frequency of 1.0%) in five total p100 samples across all timepoints. In the pDMSO populations, however, the deletion was consistently detected in all replicates and increased in frequency from a mean of 75.4% at generation 20 to a mean of 98.6% at generation 246 (**Fig. 2b-c; Table S1**). The deletion contains two putative IS3 elements and shares near-perfect identity (99.8%) with ISEc52, suggesting that the deleted variant represents the ancestral *EcN* genotype prior to IS-mediated *gnd* inactivation (**Fig. 2c**), which may have been acquired during early propagation in M9-glucose prior to treatment splitting (**Fig. 2a**).

The second variant is a missense SNP (T22I) in *sdhD*, encoding succinate dehydrogenase subunit D (**Fig. 2b; Fig. S3**). This variant uniquely persisted at ≥50% frequency in pDMSO and was undetected (<1% frequency) in all p100 replicates (**Fig. 2b; Table S1**). Although succinate dehydrogenase is amongst other membrane-bound dehydrogenases that have been reported as targets of phenothiazines^26,63^, this variant’s restriction to control populations suggests that it may be purified in TDZ adapted populations.

The remaining two variants are large (∼120 kb) deletions that were detected in p100 population replicates *b* and *c* (**Fig. 2b,d**). The two unique deletion variants share identical right-junction boundaries but differ at their left ends. Both deletions span >100 genes and include the siderophore aerobactin and the K5 CPS gene cluster, located within a previously described genomic island (“GEI-II”) (**Fig. 2d**).^50^ While only replicate *c* harbored the deletion at ≥50% frequency, we include the replicate *b* deletion as a p100-specific variant due to its similarity with replicate *c* and its elevated frequency (44.1%) by generation 246 (**Fig. 2b,d**).

We also evaluated reversible variants mediated by either DNA-flipping recombinases or slipped-strand mispairing in simple sequence repeats (SSRs).^64,65^ While programmed inversions were rare (**Fig. S4a-b**; **Table S2**), six SSRs within the ∼120-kb deletion region sustained ≥10% INDEL frequencies exclusively in the p100 populations (**Fig. S4c-e; Table S3**). Strikingly, despite lacking either CPS-deletion variant, p100 replicate *a* harbored two high-frequency SSR-associated INDELs that introduce premature stop codons in *kfiB*, a gene essential for K5 CPS surface expression (**Fig. 2e-f; Fig. S4e**).^66–68^ Functionally converging with the structural variant observed in replicates *b* and *c*, these *kfiB* loss of function variants are expected to inactivate CPS expression.

To test whether stronger selective pressure reveals missed adaptive variants, we re-sequenced generation 246 p100 populations after propagation under supra-MIC (200 µM) TDZ (**Fig. S5a; Methods**). Among variants reaching ≥10% frequency, no new mutations consistently increased in frequency between initial (i.e., “no drug”) and 200 µM TDZ conditions beyond those that were previously identified (**Fig. S5b**). Of note, the K5 CPS-affecting deletions in replicates *b* and *c* reached fixation (100% frequency) under 200 µM TDZ (**Fig. S5b**), and the *kfiB*-associated INDELs in replicate *a* increased modestly. Similarly, the ISEc52 “deletion” variant, prevalent in pDMSO controls (**Fig. 2b**), was undetected in the re-sequenced populations (**Fig. S5b**), leaving *gnd* inactivated in all p100 samples. Thus, across all three TDZ-resistant p100 replicates, selection on both O-antigen and K5 CPS loci follows consistent directional trajectories under supra-MIC drug pressure.

### KfiB-dependent CPS expression drives maximal TDZ sensitivity

The convergent loss of function of CPS-related genes in TDZ-resistant populations pointed to surface polysaccharide architecture as a potential mediator of TDZ susceptibility. This prompted us to test its causal role through targeted molecular investigations. Given that K5 CPS loss and IS-mediated *gnd* disruption were the primary variants selected under colon-approximate TDZ pressure (**Fig. 2**), we tested whether either determinant drives *EcN*’s medium-modulated susceptibility (**Fig. 1c-f**). Using an ancestral *EcN* clone as a reference (**Methods**), we generated isogenic strains with clean deletions of the *gnd*-disrupting ISEc52 element (*EcN* Δ*IS3*-*gnd*) and the CPS-essential gene *kfiB* (*EcN* Δ*kfiB*) (**Fig. 2c,e**) and then compared their drug sensitivities with that of the ancestral, wild-type clone. We did not engineer *sdhD* variants, as deletion of *sdhD* has already been shown to reduce the membrane-permeabilizing effects of TDZ.^26^

At supra-MIC (200 µM) TDZ, only Δ*kfiB* grew (**Fig. 3a**). Furthermore, at 50 and 100 µM TDZ, Δ*kfiB* exhibited markedly reduced growth inhibition compared to wild-type *EcN*, while ΔIS3*-gnd* displayed slightly increased inhibition at 100 µM (**Fig. 3a**) – both as expected. The effect of deleting *kfiB* was medium-dependent: in M9-glucose, Δ*kfiB* grew better than wild type (∼46% and ∼35% relative improvement at 50 and 100 µM TDZ, respectively), while in LB the effect was minimal (–1% and 8%, respectively) (**Fig. 3b**). To confirm that Δ*kfiB* lacks surface CPS, we separated cells via Percoll density gradients (**Fig. 3c**). Wild-type *EcN* accumulated in lighter upper fractions, whereas Δ*kfiB* consistently settled at the bottom (**Fig. 3d-e**), consistent with loss of capsule display.^69–71^ These results suggest that the K5 CPS potentiates TDZ susceptibility under nutrient limitation.

**Figure 3.**
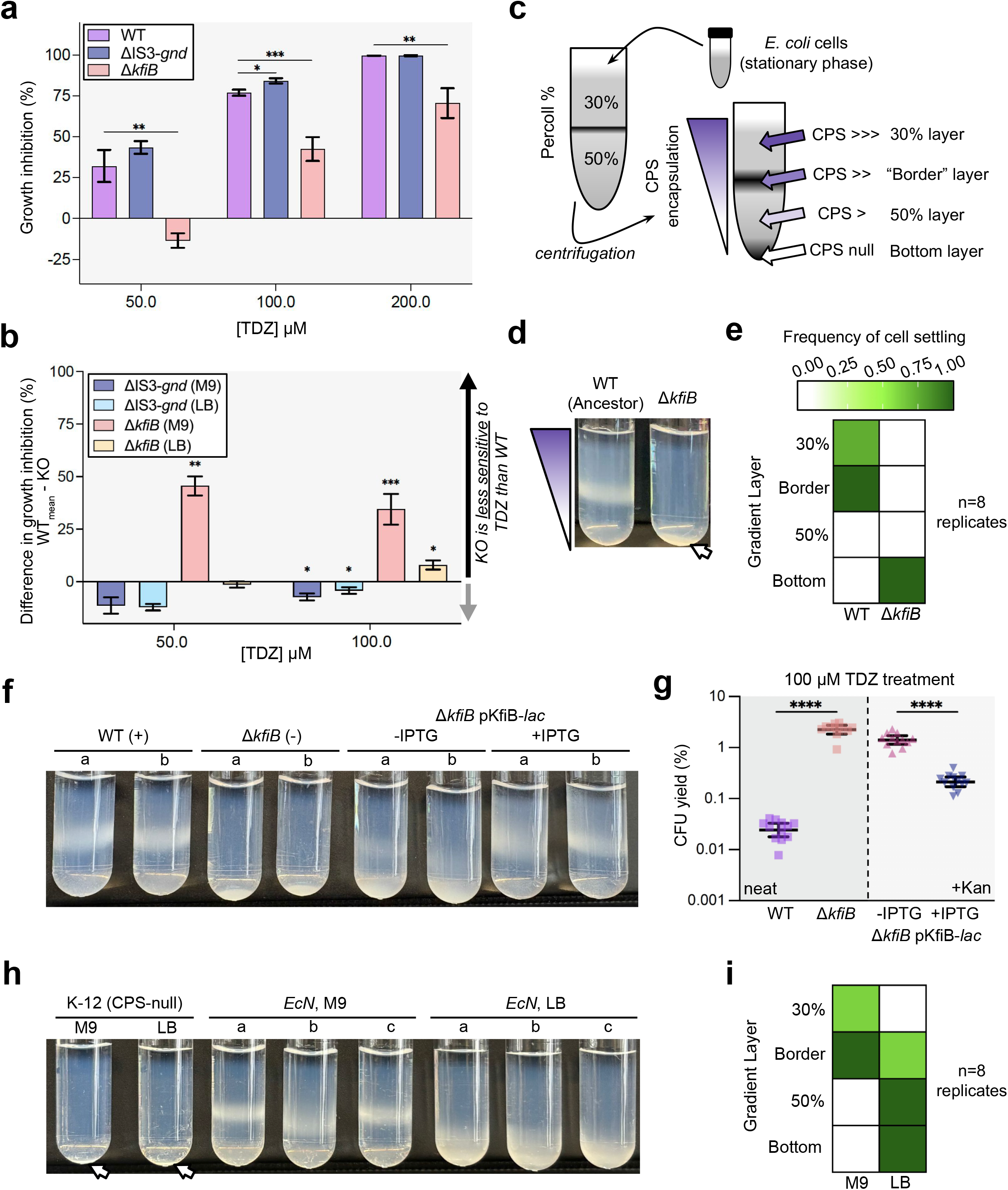
KfiB-mediated CPS display is necessary for maximal TDZ-sensitivity in minimal media. (**a**) Dose-dependent growth inhibition (%) of wild-type (WT) and single-locus knockout *EcN* clones by TDZ in M9-glucose, assessed by endpoint OD_600_ assays. Data are mean ± SEM from four biological and two technical replicates, with each knockout clone compared to WT (**b**) Difference in growth inhibition (%) between WT *EcN* and each knockout clone, tested in both M9-glucose (same data as panel A) and LB broth. LB broth growth inhibition (%) data were first gathered from two biological and two technical replicates. Data are mean ± SEM. Significance values reflect growth inhibition (%) comparisons between WT and knockout clones in each condition. (**c**) Overview of Percoll cell density gradients. CPS-expressing cells have lower density than CPS-null cells and thus settle at higher (less dense) layers in the gradient. (**d**) Representative Percoll gradient result for WT *EcN* and Δ*kfiB*. (**e**) Frequency of cell settling in specific Percoll layers for WT *EcN* and Δ*kfiB,* calculated across eight biological replicates. (**f**) Percoll gradient results for WT, Δ*kfiB*, and Δ*kfiB* pKfiB-*lac EcN*, in biological duplicate. Δ*kfiB* pKfiB-*lac* was tested without (-IPTG) and with (+IPTG) inducer (0.1 mM IPTG). (**g**) CFU yield (%) of *EcN* strains WT, Δ*kfiB*, and Δ*kfiB* pKfiB-*lac* without (-IPTG) or with (+IPTG) inducer after five hours of 100 µM TDZ treatment initiated at exponential phase. Error bars are the 95% CI centered on the geometric mean. Data are from three biological and four technical replicates. WT was compared to Δ*kfiB*, and Δ*kfiB* pKfiB-*lac* was compared between -IPTG and +IPTG. Kanamycin was included in the Δ*kfiB* pKfiB-*lac* growth medium to maintain the ectopic vector. (**h**) Representative Percoll gradient results in M9-glucose and LB broth for *EcN* in paired biological triplicates, with singlet K-12 (CPS-null) replicates as negative controls. (**i**) Frequency of cell settling in specific Percoll layers for WT *EcN* in M9-glucose and LB broth across eight biological replicates that were paired across each treatment. Mann-Whitney U was used for all comparisons: * = p < 0.05, ** = p < 0.01, *** = p < 0.001, **** = p < 0.0001. Absence of a significance value indicates a non-significant difference.

To establish causality, we reintroduced *kfiB* under an IPTG-inducible promoter on a low-copy plasmid (pKfiB-*lac*; see **Methods**) in the Δ*kfiB* strain.^72^ IPTG induction restored wild-type CPS expression levels (**Fig. 3f**) and correspondingly abolished TDZ resistance; complemented cells exhibited susceptibility similar to ancestral *EcN*, with CFU yield after 100 µM TDZ treatment significantly lower in KfiB-induced versus uninduced cells (**Fig. 3g**).

Collectively, these genetic and phenotypic experiments confirm that *kfiB*-dependent K5 CPS display is necessary for maximal TDZ susceptibility in M9-glucose, but not LB (**Fig. 3b,g**). This aligns with our observation that *EcN*’s overall susceptibility to TDZ is low in LB (**Fig. 1e-f**) and that fewer cells express surface CPS under nutrient-replete conditions (**Fig. 3h-i**). These data establish a direct mechanistic link: under nutrient limitation, K5 CPS expression paradoxically amplifies *EcN* vulnerability to TDZ.

### Small intestine-approximate TDZ exposure selects for resistant, CPS-deficient subpopulations

While 40 µM TDZ did not drive population-wide adaptation (**Fig. S2b-d**), sub-MIC antibiotic exposure is known to select for resistant subpopulations that might evade identification in bulk growth assays.^73–75^ We therefore tested whether physiologically relevant TDZ concentrations enriched for a cryptic, TDZ-resistant subpopulation. Streaking generation 20, 100, 199, and 246 glycerol stocks on 400 µM TDZ agar revealed growth of resistant colonies in p100 populations immediately by generation 20 (**Fig. S6a**). Breakthrough colonies appeared sporadically in pDMSO populations but did not increase in frequency across generations (**Fig. S6a**). Unlike either the pDMSO or p100 populations, we observed growth in p40 populations at generation 100 that increased in density by generation 246.

From the generation 246, 400 µM TDZ plates, we isolated five clones from the p100 populations and six clones (three small and three large colonies) from the p40 populations (**Fig. 4a; Fig. S6b**). OD_600_-and CFU-based growth assessments confirmed that both the p40-and p100-isolated clones had attenuated TDZ-sensitivity at physiological concentrations when compared with ancestral *EcN* and generation 246 pDMSO-isolated clones (**Fig. S6c-d**). Short-and long-read sequencing (**Methods**) revealed that four of five p100 clones carried fixed deletions within the K5 CPS locus: 100b1, 100c1, and 100c2 harbored the ∼120 kb CPS-encompassing deletions, while 100a1 harbored a fixed, single-base deletion in the second *kfiB* SSR region (**Fig. 4a; Table S4**). As would be predicted for these variants, all four were completely CPS-deficient by Percoll gradient analysis (**Fig. 4b,d**). The remaining p100 clone (100b2) had an intergenic SNP upstream of *ydiF* and exhibited a wild type-like cell density profile (**Fig. 4a-b,d**; **Table S4**), indicating alternative resistance mechanisms.

**Figure 4.**
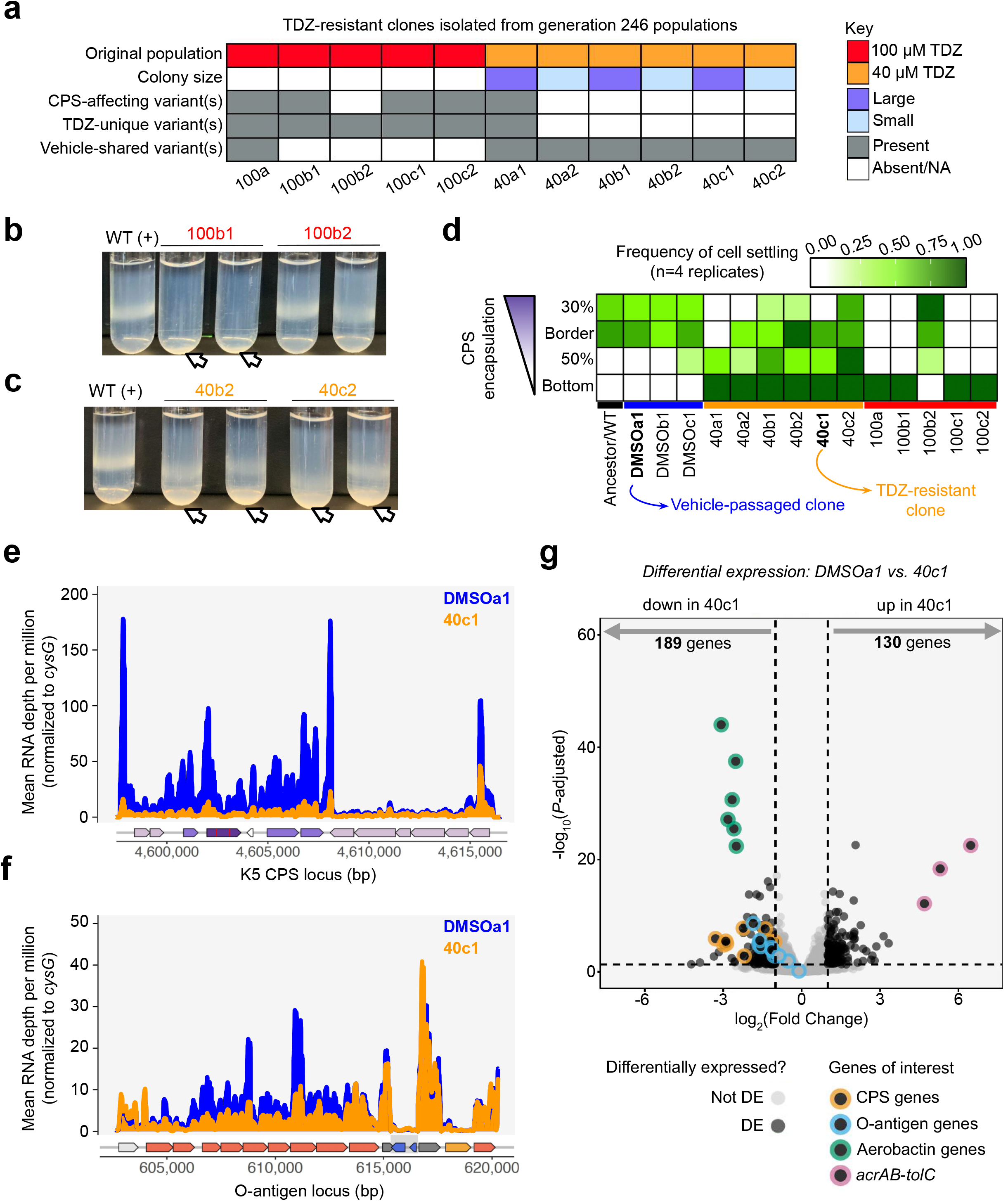
TDZ-resistant clones have reduced CPS expression regardless of DNA variants. (**a**) Summary of DNA variants identified across eleven TDZ-resistant clones isolated from generation 246 populations (see **Methods**). CPS-affecting variants are those that affect either the K5 CPS or known K5 CPS regulators. TDZ-unique variants are any variant detected in a TDZ-resistant clone that was not detected in vehicle-treated clones *or* populations, while vehicle-shared variants are those that were identified as such. (**b-c**) Representative Percoll density gradients for TDZ-resistant clones: 100b1 & 100b2 (**b**), and 40b2 & 40c2 (**c**), with WT *EcN* is included as a positive control. (**d**) Frequency of cell settling in specific Percoll layers for the eleven TDZ-resistant clones, three TDZ-susceptible clones isolated from generation 246 pDMSO populations, and WT *EcN*, each in biological quadruplicate. (**e-f**) Normalized RNA sequencing depth across the ancestral K5 CPS locus (**e**) and O-antigen locus (**f**) for the TDZ-susceptible clone DMSOa1 and the TDZ-resistant clone 40c1, averaged across biological quadruplicates. Data are from the “baseline” condition (i.e. RNA isolated after 30 minutes of vehicle treatment during exponential phase) and normalized by both library size and the housekeeping gene *cysG* (see **Methods**). (**g**) Volcano plot of differentially expressed genes between 40c1 and DMSOa1 at baseline. Differentially expressed (DE) genes fulfilled the following criteria: |log_2_(Fold Change)| ≥1.0 and *P-*adjusted <0.05, using the Benjamini-Hochberg correction for multiple hypothesis testing.

By contrast, p40-isolated clones generally lacked either CPS-affecting mutations or variants otherwise unique to TDZ-resistant *EcN* (**Fig. 4a; Table S4**). The large-colony clone 40a1 was the sole exception, carrying two fixed, CDS-truncating mutations: a transversion resulting in an early stop in the multidrug efflux regulator *acrR* and a frameshift deletion in the GTPase *bipA* (also known as *typA*).^76,77^ Although essential for growth at low temperatures, BipA also regulates maximal CPS expression at 37°C.^77,78^ Correspondingly, 40a1 settled in the bottom two Percoll layers (**Fig. 4d**), indicating reduced capsule display. Strikingly, despite lacking putative CPS-affecting mutations (**Fig. 4a**), the remaining p40 isolates displayed heterogeneous buoyant densities, with consistent subpopulations settling in the CPS-null bottom fraction (**Fig. 4c,d**).

To determine whether cell density heterogeneity reflected silencing of capsule expression in variant-free TDZ-resistant clones, we performed bulk RNA-sequencing on a representative p40 clone lacking putative resistance mutations (40c1) and a matched pDMSO control (DMSOa1), both in quadruplicate (**Fig. 4a,d**). Locus-specific RNA coverage assessments confirmed downregulation of the entire CPS locus in 40c1 compared to DMSOa1 (**Fig. 4e**) as well as reduced transcription of O-antigen glycosyltransferases (**Fig. 4f**) – both of which were significant per differential expression analysis (**Fig. 4g; Table S5**). The most prominent transcriptomic shifts included repression of the aerobactin siderophore locus (within the same genomic island as the K5 CPS; **Fig. 2d**) and marked upregulation of the *acrAB*-*tolC* efflux system, the latter having been previously associated with phenothiazine tolerance.^7,79^

Collectively, these data demonstrate that sub-MIC, small intestine-approximate TDZ exposure selects for resistant subpopulations that only emerge after strong selection (**Fig. S6a**). Despite lacking fixed CPS-affecting variants or other putative resistance mutations, p40-adapted clones have lower capsule expression (**Fig. 4a,d,g**) and higher efflux pump expression (**Fig. 4g**). Both contribute to phenotypic TDZ resistance (**Fig. 3g; Fig. S1**), likely acting in concert to alleviate vulnerability under physiologically relevant drug concentrations.

### K5 CPS potentiates the acute transcriptional response to subinhibitory TDZ

Subinhibitory (40 µM) TDZ does not significantly reduce viable cell counts in wild-type or evolved clones during standard growth assays (**Fig. S6e**). Nevertheless, because this concentration selects for resistant subpopulations (**Fig. S6a-d**), we expected susceptible cells to mount an acute transcriptional response. RNA-seq of exponential-phase DMSOa1 and 40c1 clones after 30 min of vehicle or TDZ exposure revealed distinct differentially expressed gene (DEG) profiles (**Table S5**) in response to TDZ. Consistent with phenotypic data, TDZ-sensitive DMSOa1 exhibited 458 DEGs, whereas TDZ-resistant 40c1 showed only 54 (**Fig. S7a-b**). Within DMSOa1, changes in K5 CPS, O-antigen, aerobactin, and efflux genes were modest compared to others (**Fig. S7a-b; Table S5**). The most robustly upregulated genes in DMSOa1 were *asr* and *spy* (**Fig. S7f; Table S5**), which encode periplasmic chaperones that prevent protein aggregation and promote proper folding.^80,81^ The genes *asr* and *spy* were also upregulated in 40c1 following TDZ treatment, although to a lesser magnitude (**Fig. S7f; Table S5**).

To test whether reduced K5 CPS expression drives 40c1’s muted transcriptional response to TDZ (**Fig. S7a-b**), we also assessed differential expression in wild-type *EcN* (TDZ-susceptible, like DMSOa1) and isogenic Δ*kfiB* (TDZ-resistant, like 40c1) – both “unevolved” clones. Similar to their “evolved” clone counterparts, wild type exhibited 328 DEGs upon TDZ exposure, whereas Δ*kfiB* showed only 118 (**Fig. S7c-d; Table S5**), confirming that CPS-deficient strains have a blunted response to subinhibitory TDZ.

To quantify whether the distinct backgrounds converge on a shared, CPS-driven transcriptional response, we calculated Pearson correlations between Δlog_2_(Fold Change) values across both clone comparisons: evolved (40c1 versus DMSOa1) and unevolved (Δ*kfiB* versus wild type). Using the 4,665 genes analyzed with *DESeq2* for all clones, we observed a moderate correlation (*r* = 0.44; **Fig. S7e**). However, restricting this analysis to just the genes differentially expressed in DMSOa1 increased the correlation (*r* = 0.77; **Fig. S7e**), indicating that K5 CPS expression alone may account for up to 60% of the variance in TDZ-induced transcriptional changes between DMSOa1 and 40c1. This concordance was consistent across the 16 DEGs common in all clones, which exhibited mean log_2_(Fold Change) ratios of 0.44 (median = 0.42) and 0.54 (median = 0.50) for the unevolved and evolved comparisons, respectively (**Fig. S7f**). Together, these transcriptomic analyses demonstrate that K5 CPS expression is a major determinant of *EcN*’s acute response to physiologically relevant, subinhibitory TDZ exposure.

### CPS-mediated sensitization extends to antipsychotic phenothiazines

While TDZ was explored first due to its reported antimicrobial activity against a broad selection of both gut commensals and pathobionts^7,12,26^, several other phenothiazine derivatives are clinically relevant and show potential for repurposing as antimicrobials or anticancer agents.^32,82,83^ We therefore tested whether CPS-mediated TDZ susceptibility extends across this drug class.

Using the CPS-inducible *EcN* Δ*kfiB* pKfiB-*lac* strain, we quantified CFU yields following exposure to each of five phenothiazine derivatives, along with fluoxetine (a non-phenothiazine SSRI with known antibacterial activity^59,84,85^) as a pharmacological control, at 100 µM (**Fig. 5a-c**). Induction of K5 CPS expression consistently reduced yields for all antipsychotic treatments (TDZ, chlorpromazine, trifluoperazine, and fluphenazine; **Fig. 5b**). By contrast, fluoxetine, the antihistamine promethazine, and the side chain-free parent compound exhibited no CPS-dependent susceptibility (**Fig. b-d**), indicating that sensitization is selective for specific antipsychotic derivatives rather than the broader phenothiazine chemical class.

**Figure 5.**
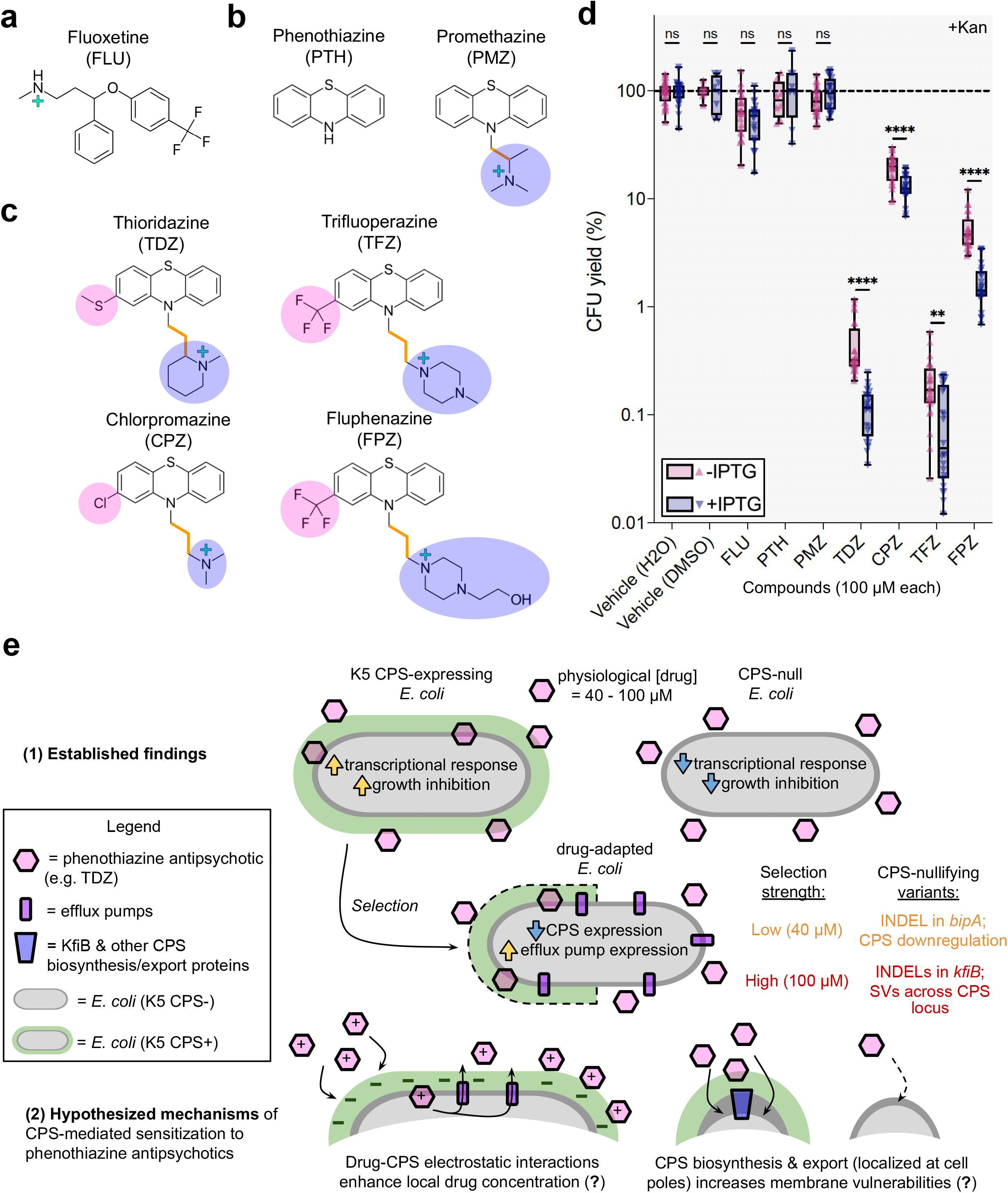
K5 CPS-mediated sensitization extends to other phenothiazine antipsychotics. (**a-c**) Compounds tested included (**a**) the SSRI fluoxetine (FLU), (**b**) non-antipsychotic phenothiazine compounds, and (**c**) phenothiazine antipsychotics including TDZ. The pink-and blue-highlighted side chains, and the number of carbons (orange) separating the phenothiazine N from the cationic N, differentiate the phenothiazines. (**d**) CFU yield (%) results for the Δ*kfiB* pKfiB-*lac* clone with (+IPTG) or without (-IPTG) induction after five hours of treatment initiated at exponential phase. Box and whiskers represent the max, min, interquartile range; the line represents the median. Data are from a minimum of two biological replicates (six for most compounds, excluding DMSO, PTH, and PMB) and four technical replicates per condition. For each compound, results were compared between -IPTG and +IPTG. Mann-Whitney U was used for all comparisons: * = p < 0.05, ** = p < 0.01, *** = p < 0.001, **** = p < 0.0001. (**e**) A model summarizing our findings and putative explanatory, mechanistic hypotheses.

The concentrations of TDZ in the human GI tract (40-100 µM) align with the threshold at which we have found selection against the K5 CPS to occur (**Fig. 5e**). This selection drives convergent capsule ablation through dose-dependent routes: 100 µM TDZ selected for structural deletions and slipped-strand mispairing-mediated INDELs affecting CPS, whereas subinhibitory 40 µM TDZ exposure predominantly selected for TDZ-resistant clones with transcriptional downregulation of CPS absent of genetic changes. Transcriptomic analyses further revealed that K5 CPS expression amplifies *EcN*’s response to subinhibitory TDZ (**Fig. S7e**). Thus, we propose that K5 CPS-mediated potentiation of TDZ’s non-antibiotic activity directly fuels selection against CPS in TDZ-adapted *E. coli* (**Fig. 5e**).

## Discussion

CPSs are generally regarded as protective structures that shield bacteria from environmental and host-derived stresses^39,41,86,87^, but our experiments suggest this view is incomplete. Rather than provide protection, the K5 CPS sensitizes *EcN* to phenothiazine antipsychotics and becomes a target of adaptive evolution under physiologically relevant drug exposure. Our findings thus support that the fitness consequences of a bacterial capsule depend on its environment: a CPS that is advantageous against one stressor can become a liability under another. This expands the growing recognition that bacterial surface architectures are shaped by multiple competing selective pressures^42,88–90^, and posits human pharmaceuticals as an additional selective force capable of remodeling bacterial surface structures. Similar exceptions to CPS-mediated “protection” have emerged in other contexts – for example, some capsules serve as phage receptors rather than barriers^90,91^, while others increase susceptibility to antimicrobial peptides^89^ – illustrating that CPS-mediated protection is not an intrinsic property of the capsule itself, but rather emerges from interactions between a particular capsule, its “host” bacterium, and specific environmental challenges, as has been previously suggested.^88^ We therefore propose that susceptibility to human drugs should be considered alongside phages and host immunity as an important determinant of capsule evolution.^42^

A major finding is that physiologically relevant phenothiazine exposure selects against capsule expression through multiple routes. Across independent evolving populations, resistance repeatedly emerged through distinct molecular mechanisms, including large chromosomal deletions, slipped-strand mispairing, and reversible transcriptional downregulation, yet converged on the same phenotypic endpoint: reduced or abolished K5 capsule expression. This convergence suggests that selection acts primarily on capsule expression rather than a single genetic pathway, highlighting the evolutionary flexibility of this trait. Experimental evolution therefore extends our genetic analyses by demonstrating that capsule loss is a readily accessible evolutionary solution to TDZ sensitivity under sustained drug exposure. The capsule loss we observed occurred through both reversible and irreversible mechanisms. The resistant clones selected under subinhibitory TDZ pressure exhibited reduced CPS expression despite lacking corresponding variants, suggesting that phenothiazine exposure can favor regulatory adaptation before stable genetic changes arise. Whether these reversible responses reflect altered nucleoid organization, DNA methylation, or selection on pre-existing phenotypic heterogeneity remains an important question for future investigation. Stronger selection, on the other hand, favored both fixed and reversible genetic alterations, with large chromosomal deletions and *kfiB*-localized slipped-strand mispairing disrupting K5 CPS expression in 100 µM TDZ-adapted *EcN*. Although slipped-strand mispairing has previously been implicated in the phase-variable regulation of capsule genes in multiple pathogens^96–98^, we are – to our knowledge – the first to suggest that it contributes to capsule regulation in *E. coli*. Together with previous observations of INDELs at the same exact locus following intestinal colonization^99^, *kfiB* appears to serve as a recurrent target of selection for modulating K5 capsule expression. These findings suggest that capsule expression is a highly evolvable trait that can respond to pharmaceutical selection through both regulatory and genetic mechanisms, allowing bacterial populations to rapidly remodel their cell surface as selective environments change.

The selective regimes we employed span drug concentrations that are expected to occur in different parts of the GI tract^14,37^. Colon-approximate (100 µM) TDZ drove rapid population-wide adaptation, whereas subinhibitory, small intestine-approximate (40 µM) TDZ more gradually enriched resistant subpopulations. These findings illustrate that clinically relevant non-antibiotic drug exposure can reshape bacterial populations long before conventional measures of growth inhibition would predict strong selection. Although the precise concentrations experienced by gut bacteria will vary with intestinal location, dose, pharmacokinetics, metabolism, and host-intrinsic factors^92–94^, our estimates were intentionally conservative (see **Methods**). Furthermore, as TDZ’s major metabolites retain pharmacological activity^37,92,95^, they may be more likely to maintain anti-CPS selection in tandem with parent compound exposure. Together, these observations support the plausibility that chronic phenothiazine treatment could impose sustained selection on capsule expression *in vivo*.

The molecular basis of K5 CPS-mediated drug sensitization remains an important question. Our findings are in line with at least two non-exclusive models (**Fig. 5e**). First, the highly anionic capsule may locally concentrate cationic phenothiazines through electrostatic interactions, increasing the effective drug concentration at the bacterial surface. Second, assembly and translocation of the capsule may create transient vulnerabilities in the cell envelope that facilitate phenothiazine entry or potentiate its membrane-disrupting effects. Although distinguishing between these possibilities will require direct biophysical measurements, both models are consistent with our observation that loss of capsule expression reduces phenothiazine susceptibility. If correct, these models suggest that bacterial capsules can actively shape the local concentration of human pharmaceuticals experienced by bacterial cells.

The evolutionary implications of our findings may extend beyond *EcN*. Previous work has shown that prolonged exposure to other antipsychotics selects for increased efflux activity^58,100^, highlighting drug-mediated evolution as an emerging feature of chronic non-antibiotic exposure. Our findings identify capsule modulation as an additional adaptive strategy. Intriguingly, transporter-dependent Group 2 *E. coli* capsules, including K5, are less prevalent among gut-associated *E. coli* than among extraintestinal infection-associated isolates.^36,101^ Although many factors undoubtedly contribute to this distribution, our findings raise the possibility that exposure to non-antibiotic drugs may represent a previously unrecognized selective pressure acting against capsule maintenance in commensal *E. coli*. Testing this hypothesis across human cohorts, animal models, and synthetic microbial communities will help determine whether pharmaceutical-driven remodeling of bacterial surface structures is a general feature of microbiome evolution.

Several limitations should be considered. First, although we demonstrate capsule-mediated sensitization across multiple phenothiazine antipsychotics, it remains unknown how broadly this phenomenon extends across the hundreds of non-antibiotic drugs that perturb the gut microbiome.^2,7^ Second, our mechanistic analyses focus on a single capsule type in a single bacterial species, whereas capsules vary enormously in polysaccharide structure, genomic architecture, and membrane translocation mechanisms across bacteria.^36,42,87^ Finally, although disruption of *kfiB* consistently abolishes capsule expression, KfiB itself participates as a chaperone of membrane-localized capsule assembly^66–68^, leaving open the possibility that some component of the observed phenotype reflects loss of KfiB-dependent membrane processes in addition to loss of the capsule. Future studies targeting other essential capsule biosynthetic genes will help distinguish these possibilities.

For decades, phenothiazine antipsychotics have been recognized as human-targeted drugs with antibacterial activity^23^, yet the bacterial determinants governing susceptibility have remained largely unknown. At the same time, CPSs have largely been viewed through the lens of host immunity, phages, and other ecological interactions.^42^ Our work links these two fields by showing that a capsule can itself determine susceptibility to non-antibiotic drugs, placing capsule expression under direct pharmaceutical selection. Although demonstrated here for a single capsule and one class of drugs, the implications may be considerably broader. Given the widespread use of human pharmaceuticals and the diversity of bacterial capsules, even modest capsule-mediated effects on drug susceptibility could have important evolutionary consequences. Medications taken daily for years or decades may therefore influence bacterial evolution not only through host physiology, but also by directly favoring or disfavoring specific surface structures. As such, recognizing medications as drivers of microbial evolution will be critical for understanding and predicting the long-term consequences of pharmacological interventions on the gut microbiome.

## Methods

### Bacterial strains

Eleven *E. coli* strains were used in this study. Seven were used without further genetic modification: wild-type Nissle 1917 (*EcN*; serotype O6:K5:K1)^50^, ATCC 31616 (strain 21; serotype O9:K35:K99), ATCC 700973 (strain C5 [Bort]; serotype O18ac:K1:H7), wild-type BW25113 and BW25113 Δ*tolC* (KEIO collection; Horizon Discovery)^102^, and the cloning strains DH5ɑ λpir and S17-1 λpir.^103,104^ According to ATCC records, ATCC 31616 was isolated from the intestinal tract of a diarrheic calf, whereas ATCC 700973 was isolated from the cerebrospinal fluid of a newborn infant.

The remaining four strains were generated in-house from a single wild-type *EcN* clone isolated from the ancestral (generation 0) population from the experimental evolution (**Fig. 2a**). These strains comprised *EcN* pSW172, *EcN* ΔIS3-*gnd*, *EcN* Δ*kfiB*, and *EcN* Δ*kfiB* pKfiB-*lac*.

### Compounds

The non-antibiotic compounds used in this study were thioridazine HCl (Millipore Sigma, T9025), chlorpromazine HCl (Millipore Sigma, C8138), phenothiazine (Millipore Sigma, P14831), promethazine HCl (Fisher Scientific, P2029), trifluoperazine 2HCl (Millipore Sigma, T8516), fluphenazine HCl USP (Fisher Scientific, NC2254207), and fluoxetine HCl (Millipore Sigma, F132). The antibiotics kanamycin sulfate USP (MedChemExpress, HY-16566A) and carbenicillin disodium salt (Fisher Scientific, ICN19509201) were used where indicated.

Powder stocks were stored at room temperature, 4°C, or −20°C according to the manufacturer’s instructions. Stock solutions were stored at −80°C unless otherwise noted; kanamycin and carbenicillin stock solutions were stored at −20°C.

### Media and culture conditions

Unless otherwise stated, bacteria were cultured aerobically at 37°C with shaking (225 rpm) in 14-mL polypropylene culture tubes (Corning, 352059). Solid cultures were incubated at 37°C on 100×15 mm Petri dishes containing 1.5% agar. Growth inhibition and growth curve assays were performed at 37°C without shaking in sterile 96-well tissue culture plates (Corning, 353072).

Two media were used throughout this study: M9-glucose and LB broth. M9-glucose contained M9 salts (3 g L^-1^ KH_2_PO_4_, 0.5 g L^-1^ NaCl, 6.8 g L^-1^ Na_2_HPO_4_, 1 g L^-1^ NH_4_Cl; Millipore Sigma, M6030), CaCl_2_ (0.011 g L^-1^; Millipore Sigma, C4901), MgSO_4_ (0.192 g L^-1^; Millipore Sigma, M7506), and D-(+)-glucose (2.40 g L^-1^; Millipore Sigma, G8270). M9-glucose plates were prepared by supplementing the same medium with 1.5% agar (Fisher Scientific, BP1423). LB broth (Fisher Scientific, BP9723) consists of casein peptone (10 g L^-1^), yeast extract (5 g L^-1^), and NaCl (10 g L^-1^), and LB plates were prepared by supplementing with 1.5% agar.

For allelic exchange, LB agar containing kanamycin (50 µg mL^-1^), carbenicillin (100 µg mL^-1^), or both was used where indicated. Counterselection was performed on sucrose agar (5% sucrose; Fisher Scientific, S5-3) prepared with Nutrient Broth (Millipore Sigma, 70122), which consists of D-(+)-glucose (1 g L^-1^), peptone (15 g L^-1^), NaCl (6 g L^-1^), and yeast extract (3 g L^-1^).

Cells were washed and serially diluted in 1X phosphate-buffered saline (PBS; 8 g L^-1^ NaCl, 0.2 g L^-1^ KCl, 1.14 g L^-1^ Na_2_HPO_4_, 0.27 g L^-1^ KH_2_PO_4;_ Fisher Scientific, AM9625) at pH 7.4.

### Comparative genomic analyses

Reference genome assemblies for *E. coli* BW25113 (GenBank: CP009273.1) and Nissle 1917 (GenBank: GCA_003546975.1) were downloaded from NCBI. Genome assemblies for ATCC 31616 (lot 70077044) and ATCC 700973 (lot 70071406) were obtained from ATCC.

Average nucleotide identity (ANI) was calculated for all pairwise genome comparisons using *FastANI* (v1.3) with default parameters.^105^ Extrachromosomal contigs were retained in all genome assemblies.

Antibiotic resistance genes were identified using the *Resistance Gene Identifier* (*RGI*; v6.0.5) from the *Comprehensive Antibiotic Resistance Database* (*CARD*; v4.0.1).^47^ Only “perfect” (100% match) and “strict” (high confidence) hits were retained.

### Estimation of GI TDZ concentrations

Estimated GI concentrations of TDZ were calculated with a previously described pharmacokinetic framework.^7^ The calculation assumes 90% absorption before the distal GI tract, small intestinal and colonic volumes of 300 mL and 600 mL, respectively, and a GI transit time of 24 hours. Using a minimum daily TDZ dose of 50 mg^37^ and an estimated fecal recovery of 50% of the administered dose^14^, estimated luminal TDZ concentrations were 41 µM in the small intestine and 102 µM in the colon. Experimental concentrations were rounded to 40 µM and 100 µM, respectively. Because the historically recommended starting dose for drug-naive individuals with schizophrenia is 50 mg three times per day^37^, these estimates are likely conservative.

### Drug sensitivity assays

Drug sensitivity was quantified using both OD_600-_and CFU-based assays. In both assays, growth in the presence of drug was normalized to a paired vehicle-treated control (≤1% DMSO) from the same biological replicate.

OD_600-_based assays were performed in 96-well tissue culture plates using geometric drug dilution series (e.g., 200, 100, 50, 25, 12.5, and 0 µM TDZ) with equivalent vehicle exposure across all conditions. Unless otherwise indicated, biological replicates (independent colonies isolated from frozen stocks) were grown overnight to stationary phase, diluted to OD_600_=0.05, and regrown to early-mid exponential phase (OD_600_∼0.3). Experimentally growing cultures were diluted 1:1000 into assay wells containing fresh medium supplemented with drug or vehicle and incubated at 37°C for 18–20 hours. Endpoint OD_600_ values were measured using an Agilent BioTek Synergy H1 microplate reader. Drug-containing blank wells were used for background subtraction. Growth inhibition (%) was calculated for each biological replicate as ((OD_600_,_vehicle_ – OD_600,drug_)/OD_600,vehicle_)×100.

To minimize potential biases arising from inoculum-dependent drug sensitivity and changes in cell size or morphology that may influence OD_600_, CFU-based assays were performed using exponentially growing cultures (OD_600_∼0.3). Cultures were divided into treatment aliquots, exposed to drug or vehicle for 5 hours at 37°C, serially diluted tenfold in sterile PBS, and spot-plated (6–8 µL) in technical replicates on drug-free LB agar. Plates were incubated overnight at room temperature before colony enumeration the following day. CFU yields were calculated for each biological replicate by normalizing drug-treated samples to the corresponding vehicle-treated control: (CFU_drug_/CFU_vehicle_)×100. Because the lowest dilution measured was 10^-1^, the theoretical limit of detection was one colony in a 6–8 µL spot, corresponding to approximately 1.3–1.7×10^3^ CFU mL^-1^.

To compare drug sensitivity between M9-glucose and LB broth, M9-glucose measurements for each strain were used as the reference condition. For OD_600_-based assays, the mean concentration-specific growth inhibition in M9-glucose was subtracted from each individual LB replicate (inhibition_LB_ - mean inhibition_M9-glucose_). For CFU-based assays, the mean CFU yield in M9-glucose was divided by the CFU yield for each individual LB replicate, and the resulting fold changes were log_2_-transformed.

To compare drug sensitivity between wild-type and single locus knockout *EcN* clones in either M9-glucose or LB broth, each replicate growth inhibition measurement for specific knockout clones was subtracted from the mean concentration-specific growth inhibition result for wild-type *EcN* (mean inhibition_wild-type_ - inhibition_knockout_).

### EcN experimental evolution

Experimental evolution was performed using a daily serial batch culture protocol similar to the *E. coli* long-term evolution experiment (LTEE).^106^ Populations were cultured for 24 hours to stationary phase before daily 1:100 transfer into fresh medium. Assuming 100-fold growth per transfer, this corresponds to approximately 6.64 generations per day and ∼246 generations after 37 transfers.^106^

A single colony of wild-type *EcN* was inoculated into M9-glucose and cultured for 24 hours to generate a common, M9-glucose-acclimated ancestral (generation 0) population. Aliquots of this population were preserved as 30% glycerol stocks and cell pellets for whole-genome sequencing.

The ancestral population was split into nine independent evolution lines (three per treatment), which were propagated in 5 mL M9-glucose cultures containing either vehicle (0.05% DMSO), 40 µM TDZ, or 100 µM TDZ. Populations were transferred daily (1:100 dilution) into treatment-containing medium for 37 consecutive transfers. Following the final treatment transfer, each population was propagated for an additional eight transfers (∼53 generations) in drug-free M9-glucose to assess the stability of evolved phenotypes after removal of selection.

Population samples were archived as 30% glycerol stocks at generations 20, 53, 100, 133, 199, and 246 using stationary-phase cultures collected immediately before each daily transfer.

### Growth curve analyses

At generations 20, 100, 199, and 246, as well as after the additional 53 generations of drug-free propagation, population-level drug sensitivity was assessed by growth curve analysis. Stationary-phase cultures were diluted 1:1000 into fresh M9-glucose containing 0, 50, 100, 200, 300, or 400 μM TDZ and transferred to 96-well plates. Drug-containing blank wells were included for background subtraction. Plates were sealed with gas-permeable membranes (Breathe-Easy; MilliporeSigma, Z380059) and incubated at 37°C in a Cerillo Stratus microplate reader, with OD_600_ measured every 10 min for 21 h.

Growth curves were analyzed in R using *gcplyr* (v.1.12.0).^60^ After subtraction of concentration-and timepoint-specific blank values, negative OD_600_ values were set to zero. For each well, the area under the growth curve (AUC), maximum culture density (OD_600, max_), and maximum growth rate over a 5-point window (Simple Derivative_max_) were calculated using *gcplyr*’s ingrained functions: auc(), max_gc(), and calc_deriv(), respectively.

### Ancestral reference genome assembly and annotation

Genomic DNA was extracted from the ancestral (generation 0) *EcN* population using the DNeasy Blood & Tissue Kit (Qiagen, 69504) according to the manufacturer’s instructions. DNA was treated with Monarch RNase A (New England Biolabs, T3018L) for 15 min at room temperature, purified using the Genomic DNA Clean & Concentrator-10 kit (Zymo Research, D4011) according to the manufacturer’s instructions, and quantified using a NanoDrop spectrophotometer.

Approximately 30 μL of DNA (≥50 ng μL^-1^) was submitted to Plasmidsaurus for whole-genome assembly using both short-read (Illumina) and long-read (Oxford Nanopore) sequencing. The resulting hybrid assembly was annotated with *Bakta* (v1.9.4, database v5.1) and used as the ancestral reference genome for all downstream variant analyses.

### Short-read genome sequencing and analysis of experimentally evolved populations

At generations 20, 53, 100, 133, 199, and 246, frozen stocks from each pDMSO and p100 experimental evolution population were revived overnight in M9-glucose medium. Genomic DNA was extracted using the same protocol described for the ancestral population. DNA samples (20 µL, ≥10 ng µL^-1^) were submitted to Novogene for paired-end Illumina sequencing (150 bp reads) using 350 bp insert libraries prepared with the ABclonal Rapid Plus DNA Library Prep Kit and sequenced on a Novaseq X Plus platform to a minimum of 2 G per sample. One sample (pDMSO, replicate *c*, generation 20) was sequenced twice to reach the minimum depth, and its resulting read files were manually concatenated prior to downstream analyses. The lowest depth sample (pDMSO, replicate *c*, generation 20) contained 13.8 million paired-end reads, with median chromosomal coverage of 393X.

Reads were quality filtered with *fastp* (v0.23.4) with default settings aside from polyG trimming (-g), paired-end overlap correction (-c), and automatic adapter detection enabled (--detect_adapter_for_pe).^107^ Processed reads were analyzed using *breseq* (v0.36.0) in polymorphism mode (-p)^62^, with the ancestral *EcN* hybrid assembly as the reference genome. Variants were detected at frequencies ≥1% (--polymorphism-frequency-cutoff 0.01), requiring a minimum of 10 reads aligned to each strand (--polymorphism-minimum-total-coverage-each-strand 10) and passing *breseq*’s strand-bias and quality-bias filters (--polymorphism-bias-cutoff 0.05).

Variants reported by *breseq* as “Predicted mutations” and “Unassigned new junction evidence” were merged in R to generate a unified variant dataset. This combined dataset included single nucleotide variants, short insertions and deletions outside of SSRs, and structural variants. Because *breseq* reports some structural variants differently depending on whether they are fixed or polymorphic, manually merging these two output tables ensured that identical structural variants were represented as single events in downstream analyses. Unless otherwise indicated, downstream analyses included variants reaching ≥3% frequency in at least one sample. For focused analyses of adaptive evolution, *breseq*-detected variants reaching ≥50% frequency in at least one replicate within only one treatment group were prioritized for further analysis. The single exception is the 44.1% frequency (at generation 246) deletion in p100 replicate *b* that is near-identical to a ≥50% frequency deletion in p100 replicate *c*.

Structural visualization of the pDMSO-associated SdhD T22 residue was performed in *ChimeraX* (v1.11.1) using an *E. coli* succinate dehydrogenase crystal structure (PDB: 1NEN).^108^

Genome-wide sequencing coverage was calculated using *breseq* BAM2COV with single-nucleotide resolution (--table --resolution 0 --region contig_1:1-5056684), including “total” coverage (i.e., coverage contributed by both uniquely mapped and multi-mapped reads).

The insertion sequence associated with the recurrent *gnd* deletion was identified by querying the deleted genomic interval against *ISFinder* using its server-ingrained *BLASTn* tool.^109^ The highest-scoring match corresponded to ISEc52 (bit score = 2478; E-value = 0), with complete sequence identity except for three nucleotides outside the annotated element boundaries, hence 99.8% total identity.

Recombinase-mediated invertible DNA elements (invertons) were identified using *PhaVa* (v0.2.3).^110^ Candidate inverted repeat-flanked regions were first identified in the ancestral reference genome, after which reads from each evolved population were aligned to reference and inverted sequence databases using *Bowtie2* (v2.4.1).^111^ Inverton frequencies were calculated using *PhaVa*’s short-read algorithm, considering only junction-spanning reads to minimize false-positive inversion calls.

To identify slipped-strand mispairing (SSM) events, custom *Python* (v3.7.16) scripts were used to identify homopolymeric SSRs within the ancestral genome and quantify SSR-associated 1-2 bp insertions and deletions from aligned sequencing reads. Analyses were restricted to SSRs ≥6 bp in length because mutation frequency increases strongly with repeat length^97,112^ and shorter repeats (>200,000 loci) substantially increase background signal. SSM frequencies were calculated as the fraction of SSR-spanning reads containing qualifying 1-2 bp SSR-consistent INDELs relative to all informative reads spanning that SSR. Specifically, the reads containing 1-2 bp INDELs strictly within the SSR region were counted (“SSM-INDEL” reads), requiring same-nucleotide identity for insertions. To capture potential SSM-mediated insertions that add tailing nucleotides outside of the original SSR boundary, 1-2 bp insertions of SSR-identical nucleotides after the right-end coordinate were also counted in the SSM-INDEL group. Because SSM-mediated INDELs are generally of the same identity as the simplest repeating unit of the given SSR^97,112^, SSR-localized INDELs ≥3 bp were measured but *not* included in either the “reference” read group (i.e. SSR-aligning reads that don’t have a qualifying INDEL) or the SSM-INDEL read group, as they represent non-SSM variation. For the same reason, reads that included ≥1 bp insertions of *non*-SSR nucleotides, and reads that included SSR deletions that also extended into neighboring non-SSR nucleotides, were each measured but not included in either read group. The per-population and per-timepoint frequencies of 1-2 bp SSM-mediated INDELs were thus calculated from SSR-aligning reads as such: Reads_SSM-INDELs_/(Reads_reference_ + Reads_SSM-INDELs_). Putative *kfiB* SSM events were manually inspected using *IGV* (v2.13.0).^113^

### Re-sequencing of populations grown in 200 µM TDZ

For each generation 246 p100 population, frozen stocks were revived overnight in M9-glucose containing 200 µM TDZ. DNA extraction, library preparation, Illumina sequencing, read processing, and variant calling were performed as described for the originally sequenced populations.

Variant frequency comparisons between the original (“no drug”) generation 246 populations and the 200 µM TDZ re-sequenced populations were restricted to (i) variants reaching ≥10% frequency following growth in 200 µM TDZ and (ii) variants prioritized in the longitudinal sequencing analysis, including recurrent capsule-associated structural variants, putative SSM events in *kfiB*, the *sdhD* T22I substitution, and the *gnd*-restoring deletion.

### Genetic engineering of EcN

Markerless chromosomal deletions and plasmid complementation strains were generated from the ancestral *EcN* isolate using allelic exchange and plasmid transformation. Allelic exchange was performed using the *sacB*-containing suicide vector pGP706 as previously described.^114^ A single clone isolated from the ancestral (generation 0) *EcN* population was selected as the parental strain for engineering. Hybrid whole-genome sequencing (Plasmidsaurus) confirmed near identity to the ancestral reference genome, differing only by two intergenic 1-bp deletions located >100 bp from the nearest coding sequence, while retaining the ISEc52 insertion within *gnd*. The heat-sensitive, carbenicillin-resistant plasmid pSW172 was introduced by electroporation to generate the parental engineering strain *EcN* pSW172.

Deletion constructs targeting *kfiB* and the IS3 insertion within *gnd* were assembled in SphI-digested pGP706 using PCR-amplified homology arms (∼500 bp flanking each side of the intended deletion) and the NEBuilder HiFi DNA Assembly Master Mix (NEB, E2621). Plasmids (pGP706-Δ*kfiB* and pGP706-ΔIS3-*gnd*) were maintained in *E. coli* DH5α λpir, verified by complete plasmid sequencing (Plasmidsaurus), and subsequently transferred into the conjugation donor strain S17-1 λpir. Conjugation with *EcN* pSW172 and allelic exchange were performed as previously described.^114^ Briefly, exconjugants were selected on kanamycin-and carbenicillin-containing LB agar, followed by sucrose counterselection to resolve the second homologous recombination event and generate the markerless deletion strains *EcN* Δ*kfiB* and *EcN* ΔIS3-*gnd*. Whole-genome sequencing by Plasmidsaurus confirmed the expected clean deletions in both engineered strains.

To generate the complementation plasmid, the low-copy plasmid pWSK129^72^ was modified by cloning the wild-type *kfiB* coding sequence together with its upstream intergenic region downstream of the endogenous *lac* promoter using NEBuilder HiFi DNA Assembly (NEB, E5520S). The resulting plasmid (p*KfiB*-lac) was verified by Sanger sequencing (Elim Biopharmaceuticals) and introduced into electrocompetent *EcN* Δ*kfiB* by electroporation, generating the strain *EcN* Δ*kfiB* p*KfiB*-lac. Transformants were maintained under kanamycin selection.

### Percoll density gradients

Cell density gradients were performed using Percoll (Millipore Sigma, P4937) as previously described.^69–71^ Individual biological replicates (independent colonies) were grown overnight (16-18 hours) in 5 mL of drug-free M9-glucose to stationary phase. Cultures prepared on the same day were normalized to equivalent OD_600_, pelleted, resuspended in 500 µL of PBS, and layered onto preformed Percoll gradients consisting of 2 mL of 30% Percoll over 2 mL of 50% Percoll (with Percoll diluted in PBS).

Gradients were centrifuged at 3,000*×*g for 30 min at 10°C using a 5920 R centrifuge (Eppendorf, 5948000131) with low acceleration (setting 3) and no braking (setting 0) to preserve gradient integrity. Following centrifugation, the presence or absence of cells was recorded for each of four density fractions: the pellet (below the 50% Percoll layer), the 50% Percoll layer, the interface between the 30% and 50% Percoll layers, and the 30% Percoll layer. Gradients were imaged using a smartphone camera. For each strain, the frequency of cell settling within each density fraction was calculated across a minimum of four biological replicates as (Replicates with cells in layer)/(Total biological replicates).

### KfiB complementation assays

Growth inhibition assays and Percoll density gradient centrifugation were performed for the complementation strain *EcN* Δ*kfiB* pKfiB-*lac* as described above. Kanamycin (50 µg mL^-1^) was included throughout all experiments to maintain plasmid selection. Expression of *kfiB* was induced with 0.1 mM IPTG (MilliporeSigma, I6758), which was added either to overnight cultures for Percoll density gradient experiments or to OD_600_-normalized subcultures (OD_600_=0.05) prior to CFU-based drug sensitivity assays.

### Isolation of TDZ-resistant and TDZ-sensitive clones

TDZ-resistant clones were isolated from experimentally evolved populations at generations 20, 100, 199, and 246 by plating frozen population samples directly onto M9-glucose agar containing 400 μM TDZ. Plates were incubated at 37°C for 48 hours. For downstream analyses, clones were isolated from the generation 246 p40 and p100 populations. Each selected colony was again streaked on fresh 400 μM TDZ plates to ensure clonality. Because colony size heterogeneity was readily apparent among p40 populations, one small and one large colony were intentionally isolated from each p40 replicate. By contrast, colony size was uniform among p100 populations; one colony was isolated from replicate *a* and two colonies each from replicates *b* and *c*. Restreaked colonies were cultured overnight in 2 mL of M9-glucose containing 200 μM TDZ before preparation of 30% glycerol stocks and cell pellets for downstream DNA extraction. Original 400 μM TDZ selection plates were imaged after 72 hours using an Epson Perfection V600 Photo scanner, with contrast modified using *ImageJ 2* (v1.54i).

TDZ-sensitive control clones were isolated from generation 246 pDMSO populations by plating frozen samples onto drug-free M9-glucose agar. Two colonies were isolated from each biological replicate, restreaked to ensure clonality, cultured overnight in drug-free M9-glucose, and stored as 30% glycerol stocks and cell pellets for downstream DNA extraction.

### Sensitivity profiling of clones isolated from experimentally evolved populations

A total of 17 experimental evolution-derived clones were characterized using both OD_600_-and CFU-based drug sensitivity assays, including six clones isolated from pDMSO populations (DMSOa1/a2, DMSOb1/b2, DMSOc1/c2), six clones isolated from p40 populations (40a1/a2, 40b1/b2, 40c1/c2), and five clones isolated from p100 populations (100a, 100b1/b2, 100c1/c2). Drug sensitivity was compared with that of ancestral *EcN* isolated from the generation 0 population. OD_600_-and CFU-based assays were performed as described above.

To preserve the phenotypic state under which resistant clones had been selected, p40-and p100-derived isolates were maintained under continuous TDZ exposure before sensitivity testing. Briefly, these clones were recovered from glycerol stocks on M9-glucose agar containing 400 μM TDZ, cultured overnight (in biological replicates) in M9-glucose containing 200 μM TDZ, washed twice with PBS to remove residual drug, and then used to inoculate drug-free subcultures (OD_600_ = 0.05) immediately before sensitivity assays. By contrast, ancestral *EcN* and pDMSO-derived clones were recovered and cultured entirely in drug-free M9-glucose.

To assess the stability of evolved drug resistance in the absence of continued selection, a subset of resistant isolates (40a1, 40b2, 40c2, 100a, 100b2, 100c1) were re-profiled after recovery and overnight growth in drug-free M9-glucose. Drug sensitivity assays were then performed as described above and analyzed together with the initial experiments, with ancestral *EcN* included in both batches as a common TDZ-sensitive control. Clone 40a1 produced colonies that were too small for reliable CFU enumeration in the repeat experiment and was therefore excluded from the repeated analysis.

### Short-read DNA sequencing and analyses of TDZ-resistant EcN clones

Following confirmation of their TDZ-resistant phenotypes, genomic DNA was extracted from the previously archived p40-and p100-derived clone pellets using the same protocol described for the experimentally evolved populations. Whole-genome sequencing was performed by Novogene using paired-end Illumina sequencing to a minimum of 1 G per sample. The lowest-depth sample (40a1) contained 9.5 million paired-end reads, with median chromosomal coverage of 274X.

Read processing, variant calling, and all downstream genomic analyses were performed as described for the experimentally evolved populations, including *breseq* polymorphism analysis, *PhaVa*-based inverton analysis, and custom SSM detection.

### Long-read DNA sequencing and analysis of TDZ-resistant and TDZ-susceptible EcN clones

All 11 p40-and p100-derived clones previously characterized by short-read sequencing were recovered from glycerol stocks on M9-glucose agar plates containing 400 µM TDZ. Single colonies were cultured overnight in 2 mL of M9-glucose containing 200 µM TDZ, and genomic DNA was extracted using the same protocol described for short-read sequencing. Genomic DNA was also prepared from three pDMSO-derived clones (DMSOa1, DMSOb1, DMSOc1). Libraries were prepared from 400 ng of DNA using the Oxford Nanopore Native Barcoding Kit 96 V14 (SQK-NBD114.96) and sequenced on a PromethION platform.

Raw signal (.pod5) files were basecalled, demultiplexed, and adapter trimmed using *Dorado* (v0.9.1; dna_r10.4.1_e8.2_400bps_sup@v5.0.0 model). Reads were converted to FASTQ format using *samtools* (v1.23.1)^115^ and filtered with *NanoFilt* (v2.8.0)^116^ to retain reads ≥1 kb and Q≥10. Following filtering, the median read quality was Q=16.6, the median read N50 was 9.7 kb, and the lowest-coverage sample achieved approximately 70X genome coverage, as estimated using *NanoPlot* (v1.41.6).^117^

Filtered reads were aligned to the ancestral *EcN* reference genome using *ngmlr* (v0.2.7; -x ont)^118^, followed by BAM conversion, sorting, and indexing with *samtools*. Single nucleotide variants and small INDELs were identified using *clair3* (v1.1.2)^119^ with parameters optimized for haploid bacterial genomes (--platform ont --include_all_ctgs --haploid_precise --no_phasing_for_fa --enable_long_indel) and the ONT Rerio model corresponding to the *Dorado* basecalling model (dna_r10.4.1_e8.2_400bps_sup@v5.0.0).^120^ Only variants with *clair3* quality scores ≥Q10 and allele frequencies ≥50% were retained.

Structural variants ≥50 bp were identified using *Sniffles2* (v2.5.3).^121^ Reads with excessive split alignments were excluded using a dynamic split-read filter (--max-splits-base 10 --max-splits-kb 0.5). Only structural variants present at frequencies ≥50% were retained for downstream analyses; all retained variants had *Sniffles2* quality scores >Q50.

Invertible DNA elements were analyzed using *PhaVa* (v0.2.3) in long-read (i.e., default) mode. No invertons were supported by more than one read in the inverted orientation, and only two loci exhibited a single inversion-supporting read. SSM analyses were not performed on long-read data because homopolymer-associated insertion and deletion errors remain enriched in Oxford Nanopore sequencing.

Complete genome assemblies were generated from filtered Nanopore reads using *Flye* (v2.9.4) with the --nano-raw and --keep-haplotypes options and otherwise default parameters.^122^

### Classification of fixed variants in EcN clones

“Fixed” variants were identified in TDZ-resistant clones with relaxed frequency thresholds to capture clonal heterogeneity that may contribute to observed phenotypes. Variants were initially selected from the *breseq* results if they had ≥50% frequency, or from the *PhaVa* and SSM-INDEL analysis results if they had ≥25% frequency. *Clair3*-, *Sniffles2*-, and *PhaVa* long-read variant analyses - as well as *Flye*-assembled chromosomes - were then used to compare and confirm the short-read-detected variants. SNPs and ≤50 bp INDELs were considered confirmed by *clair3* if they exceeded 50% frequency; SVs >50 bp were considered confirmed by either *Sniffles2* if they exceeded 50% frequency or by *Flye* if they were present in the assembled chromosome; inversions were considered confirmed if they exceeded 25% frequency in the long-read *PhaVa* analysis. There were no SVs or inversions detected by *Sniffles2* or long-read *PhaVa* (within the above parameters) that were not also detected with short-read analyses.

In TDZ-susceptible, pDMSO-isolated clones, fixed variants were identified with *clair3*, *Sniffles2*, and long-read *PhaVa*. Variants detected by either *clair3* or *Sniffles2* at ≥50% frequency, or *PhaVa* at ≥25% frequency, were considered fixed.

Finally, each of the fixed variants was classified as one (or multiple) of the following: “vehicle-shared,” “TDZ-unique,” or “CPS-affecting.” Vehicle-shared variants were those that were detected as fixed variants in at least one pDMSO-isolated clone, or that were identified at ≥10% frequency in at least one pDMSO replicate population at any generation. Unique variants otherwise predicted to have the same molecular consequence (e.g., early stop mutations in the same gene) were considered “shared.” TDZ-unique variants were those variants that were only detected in p40-and p100-isolated clones (and populations). CPS-affecting variants were those with predicted molecular consequences affecting the K5 CPS locus or known K5 CPS-affecting regulators (e.g., *bipA*). All CPS-affecting variants were also TDZ-unique variants.

### RNA isolation, sequencing, and analysis

Total RNA was isolated from four *EcN* genotypes: the experimentally evolved TDZ-resistant clone 40c1, the TDZ-sensitive clone DMSOa1, the parental engineering strain *EcN* pSW172, and *EcN* Δ*kfiB*. *EcN* pSW172 was cultured at 37°C before experimentation to cure the temperature-sensitive pSW172 plasmid, thereby generating a wild-type background directly comparable to *EcN* Δ*kfiB*. Each genotype was analyzed in biological quadruplicate across two independent experimental batches (40c1 and DMSOa1 in batch 1; *EcN* pSW172 and *EcN* Δ*kfiB* in batch 2).

Cultures were recovered from glycerol stocks on drug-free LB agar, grown overnight in M9-glucose, diluted to OD_600_ = 0.05, and regrown to mid-exponential phase (OD_600_∼0.25). Each biological replicate was then split into two treatment conditions: 40 μM TDZ or vehicle (0.1% DMSO) for 30 min. Vehicle-treated samples were considered representative of “baseline” transcriptomes for downstream analyses. Cultures were immediately quenched with ice-cold acidic phenol/ethanol and total RNA was extracted using a previously described protocol and the Quick-RNA Fungal/Bacterial Miniprep Kit (Zymo Research, R2014).^110^ Following DNase treatment (Turbo DNase, AM2238; Invitrogen), RNA was purified using the RNA Clean & Concentrator-5 kit (Zymo Research, R1014). RNA concentration was measured using the Qubit RNA HS assay and purity assessed by NanoDrop spectrophotometry. From 32 samples, libraries were prepared and sequenced by SeqCoast Genomics using the Illumina Stranded Total RNA Prep with Ligation, Ribo-Zero Plus Microbiome Kit (Illumina, 20072063) on an Illumina NextSeq 2000 platform (150 bp paired-end reads). Mean RNA integrity (RIN) across all samples was 7.9.

Reads were trimmed and quality filtered using *fastp* (v0.23.4), retaining reads ≥25 nt with an average quality score ≥Q20. Reads were aligned to the ancestral *EcN* reference genome using *Bowtie2* (v2.5.4) with sensitive local alignment, and transcript abundance was quantified using *featureCounts* (v2.0.8)^123^ with paired-end counting and reverse-strand specificity.

To quantify transcription across the O-antigen and CPS loci, per-base sequencing coverage was calculated using *samtools* (v1.23.1) and normalized in R with *dplyr* (v1.2.0) (dplyr.tidyverse.org). Coverage at each genomic position was first normalized to total mapped library size and then divided by the normalized mean coverage of the housekeeping gene *cysG*, which was selected because it was stably expressed across all experimental conditions and has previously been validated as a reference gene.^124,125^ Final values are reported as mean normalized reads per million (RPM) across biological quadruplicates.

Differential expression analysis was performed using *DESeq2* following median-of-ratios normalization. Genes with fewer than 10 total reads across all samples and rRNA features were excluded. Differentially expressed genes were defined as those with |log_2_(Fold Change)| >1 and Benjamini–Hochberg adjusted *P*<0.05.

To compare transcriptional responses between independently evolved and genetically engineered TDZ-resistant strains, Pearson correlations were calculated in R using TDZ treatment-induced Δlog_2_(Fold Change) values for 40c1 versus DMSOa1 and Δ*kfiB* versus wild-type *EcN*. Correlations were calculated using either all expressed genes retained by *DESeq2* (4,665 genes) or only genes that were differentially expressed in DMSOa1 (452 genes). While there were 458 DEGs in DMSOa1, six were not used for the correlation analysis due to their absence in the *DEseq2* output for either wild type, Δ*kfiB*, or 40c1.

### Statistics

Statistical analyses were performed using GraphPad Prism and R. Unless otherwise indicated, statistical comparisons were performed using two-sided Mann-Whitney U tests. Statistical tests both sample size and *P* value ranges are provided in the corresponding figure legends. Exact sample sizes, including biological replicates, and *P* values are provided in the supplement (see **Table S6**, and **Tables S5 & S7**, respectively).

### Figure generation

Graphs were generated using GraphPad Prism and R. Multi-panel figures were assembled and annotated using Google Drawings and Affinity Designer.

## Data and code availability

All DNA and RNA sequencing data, together with the wild-type (ancestral and pSW172) *EcN* genome assemblies, have been deposited in the NCBI Sequence Read Archive (or GenBank, respectively) under BioProject accession PRJNA#######. The custom scripts used to detect SSRs in the *EcN* reference genome and calculate their putative SSM-mediated 1-2 bp INDEL frequencies have been made accessible on GitHub (https://github.com/moogill/TDZ-CPS-Project).

## Acknowledgements

This work was supported in part by the G. Harold & Leila Y. Mathers Foundation (MF-2112-02177 awarded to E.T.K. and A.S.B.) and the NSF Graduate Research Fellowship Program (awarded to M.O.G.). H.S. acknowledges support from the James S. McDonnell Postdoctoral Fellowship, W.L. acknowledges support from the Knut and Alice Wallenberg Foundation Bienenstock Postdoctoral Fellowship, and R.B.C. acknowledges support from the A.P. Giannini Postdoctoral Research Fellowship. E.T.K. acknowledges the NIH (GM145357), and L.C. acknowledges the National Institute of General Medical Sciences of the NIH (R35GM156332), for additional support. The Bhatt lab is supported by NIH U54AG089334, NIH U01DE035635, NIH R01CA301727 and NHLBI R01HL181969.

We thank Dr. Jakob Wirbel and Dr. Jordan Lin for their preliminary readings of the manuscript. We also thank Dr. Andrew Verdegaal, Dr. Eitan Yaffe, Dr. Soumaya Zlitni, Dr. Ivan Zheludev, Dr. Michael J. McDonald, and members of the Huang and Bhatt laboratories for invaluable discussions and feedback on the manuscript.

## Author contributions

M.O.G. and A.S.B. designed the study. Experiments and data processing were performed by M.O.G., J.A.C., K.X.J., A.A., H.S., A.N., R.B.C., Z.B., W.L., and D.T.S. Data interpretation was performed by M.O.G., J.A.C., K.X.J., and A.S.B. Data visualization was performed by M.O.G., K.X.J., and R.B.C. Funding was procured by A.S.B., E.T.K., K.C.H., and L.C. The original draft was written by M.O.G. and A.S.B. Subsequent drafts were edited by M.O.G., J.A.C., K.X.J., A.N., R.B.C., D.T.S., E.T.K., G.S., K.C.H., and A.S.B. All authors reviewed the manuscript.

## Supplementary Data

**Supplementary Table 1. Frequencies of *breseq*-detected variants in experimentally evolved populations.**

**Supplementary Table 2. Inversion frequencies of PhaVa-detected invertons in experimentally evolved populations.**

**Supplementary Table 3. Frequencies of 1-2 bp INDELs across ≥6 bp SSRs in experimentally evolved populations.**

**Supplementary Table 4. Fixed variants in generation 246-isolated clones.**

**Supplementary Table 5. DEseq2 results for the DMSOa1-vs-40c1 clone comparison and all clone-specific TDZ-vs-vehicle comparisons.**

**Supplementary Table 6. Summary of sample sizes and replicates across experiments.**

**Supplementary Table 7. Summary of *P*-values for all two-sided Mann-Whitney U comparisons.**

**Supplementary Figure 1.**
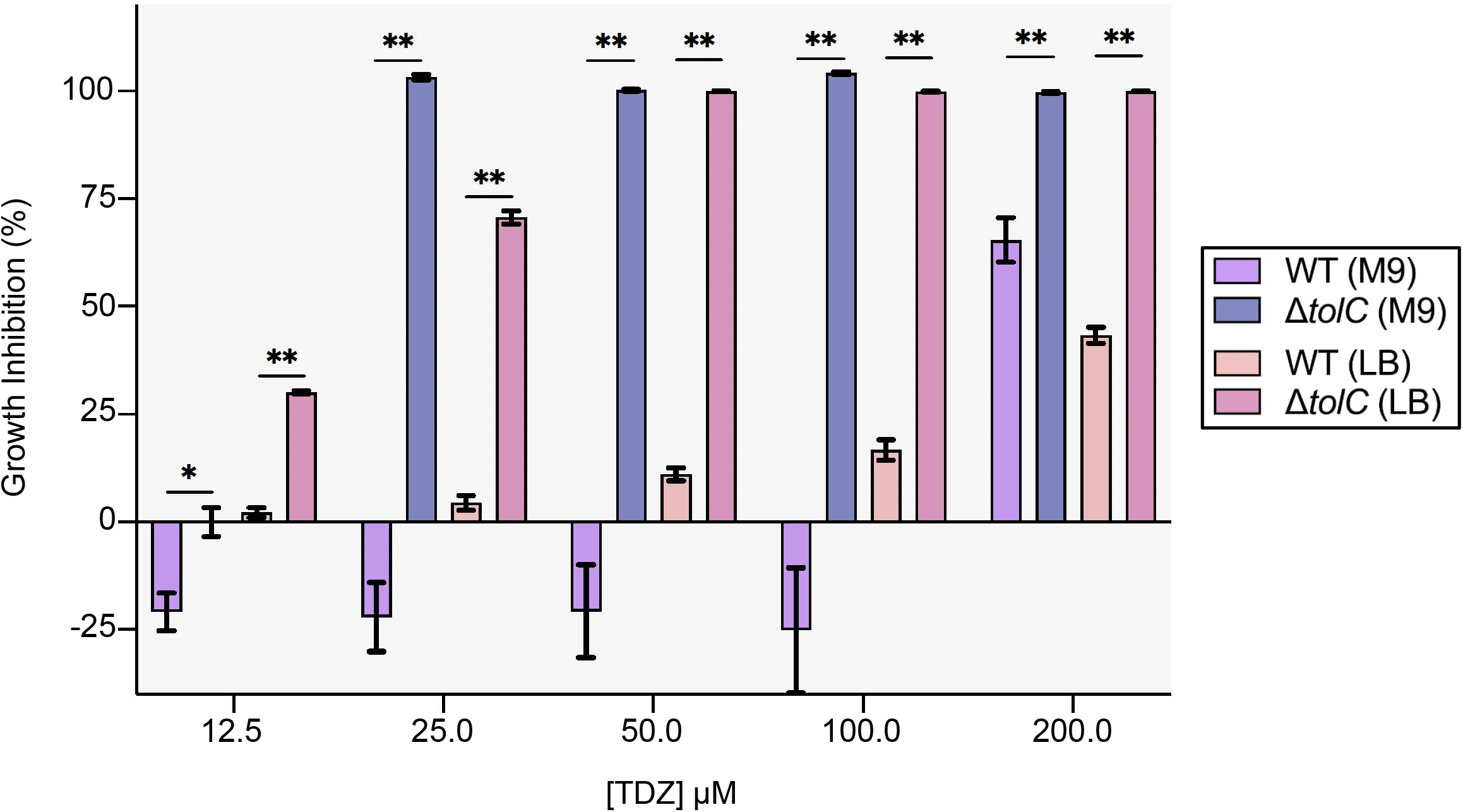
Resistance to phenothiazine antipsychotics is mediated by *tolC* in *E. coli*. Dose-dependent growth inhibition (%) of WT and Δ*tolC* K-12 BW25113 by TDZ in M9-glucose and LB broth, assessed by endpoint OD_600_ assays. Data are mean ± SEM from two biological and three technical replicates. Within each medium, WT was compared to Δ*tolC* K-12. Mann-Whitney U was used for comparisons: * = p < 0.05, ** = p < 0.01, *** = p < 0.001, **** = p < 0.0001.

**Supplementary Figure 2.**
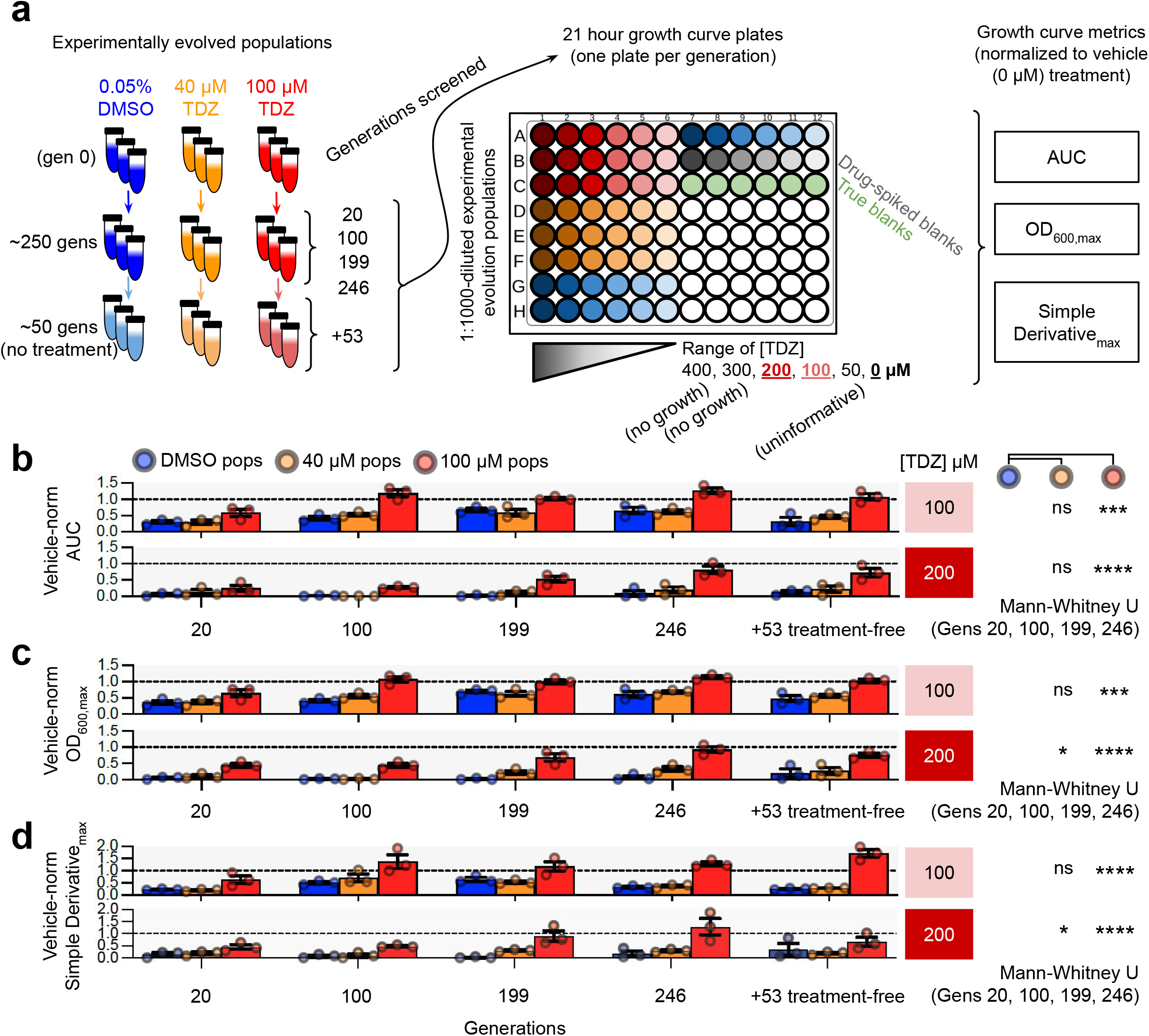
Colon-approximate TDZ-exposure selects for population-level resistance. (**a**) Overview of the growth curve assays used to assess TDZ-adaptation in the nine experimentally evolved populations across four treatment timepoints (generations 20, 100, 199, 246), and after 53 generations without treatment. (**b-d**) Results of the (**b**) AUC, (**c**) OD_600,max_, and (**d**) Simple Derivative_max_ measurements for each treatment group assessed across generations. Data are mean ± SEM after normalization by the mean results for the vehicle (i.e. “0 µM”) controls at each generation; the dashed line (y=1) represents equivalency with vehicle treatment. Mann-Whitney U was used to compare the p40 and p100 populations results, across generations 20, 100, 199, and 246, to the pDMSO populations: * = p < 0.05, ** = p < 0.01, *** = p < 0.001, **** = p < 0.0001.

**Supplementary Figure 3.**
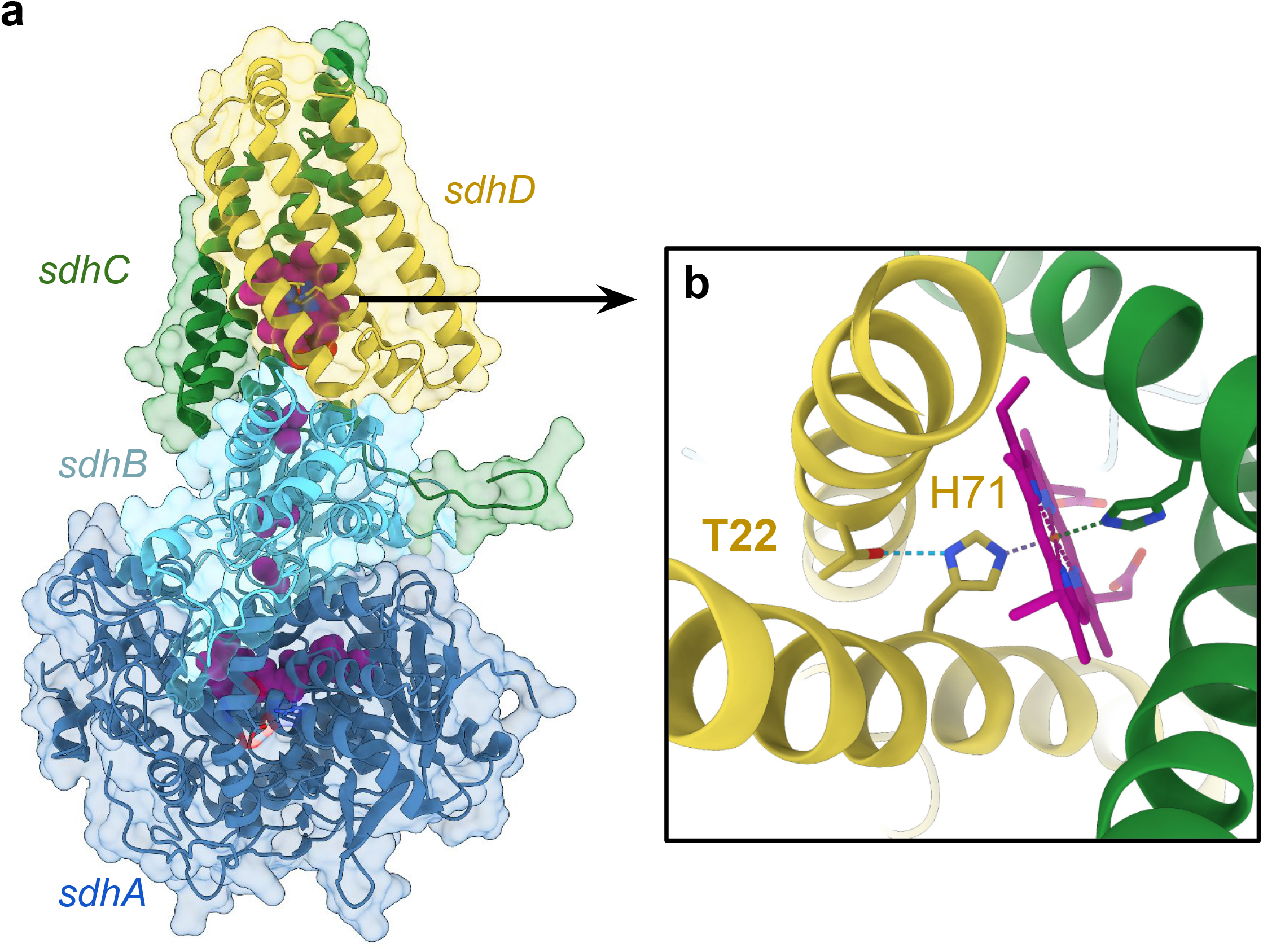
Threonine-22 forms a hydrogen bond with heme-coordinating Histamine-71 in *sdhD*. (**a**) Crystal structure of an *E. coli* succinate dehydrogenase complex (PDB: 1NEN), with the *sdhD* subunit highlighted in yellow. (**b**) Zoomed-in view of Threonine-22 and its H-bond with Histidine-71 in *sdhD*. Histidine-71 also forms an H-bond with heme.

**Supplementary Figure 4.**
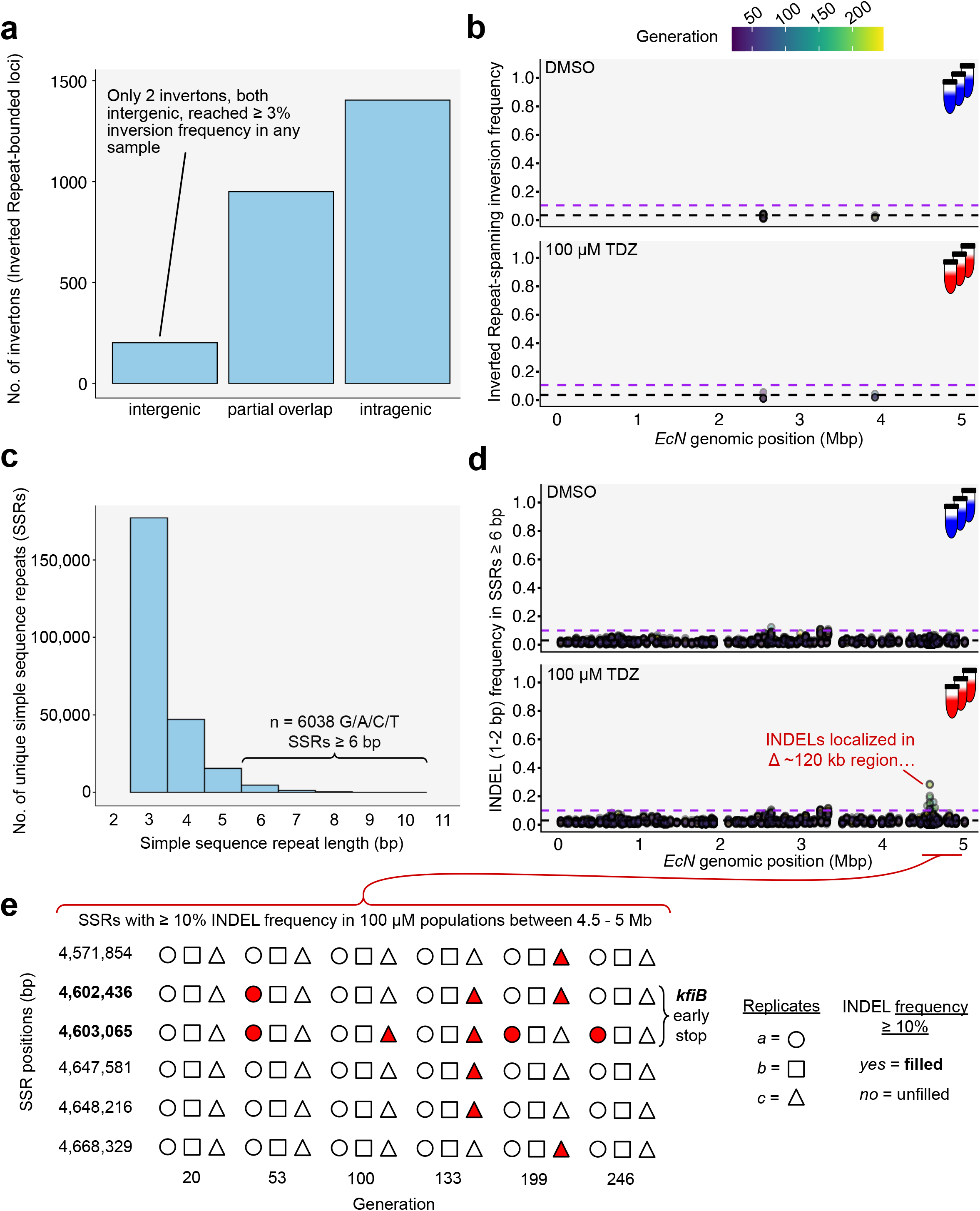
*KfiB*-localized INDELs are the sole reversible DNA variants to differentiate pDMSO and p100 populations. (**a**) The number of *PhaVa*-detected putative invertons (i.e. inverted repeat-bounded loci) in the ancestral *EcN* reference genome, categorized by whether they are intergenic or located either entirely (“intragenic”) or partially (“partial overlap”) within a gene. (**b**) Inversion frequencies of those invertons reaching ≥3% (black dashed line) in at least one replicate for any population. No invertons ever reached ≥10% (purple dashed line) inversion frequency. (**c**) The number of simple sequence repeats (SSRs) in the ancestral *EcN* reference genome, categorized by total SSR length (bp). No SSRs ≥11 bp were detected, and 2 bp repeats were ignored. (**d**) Frequencies of 1-2 bp INDELs in ≥6 bp SSRs (see **Methods**) in the pDMSO and p100 populations across generations. (**e**) Summary of the generation-specific, p100 population replicates that harbored SSRs with ≥10% INDELs between 4,500,000 bp and 5,000,000 bp.

**Supplementary Figure 5.**
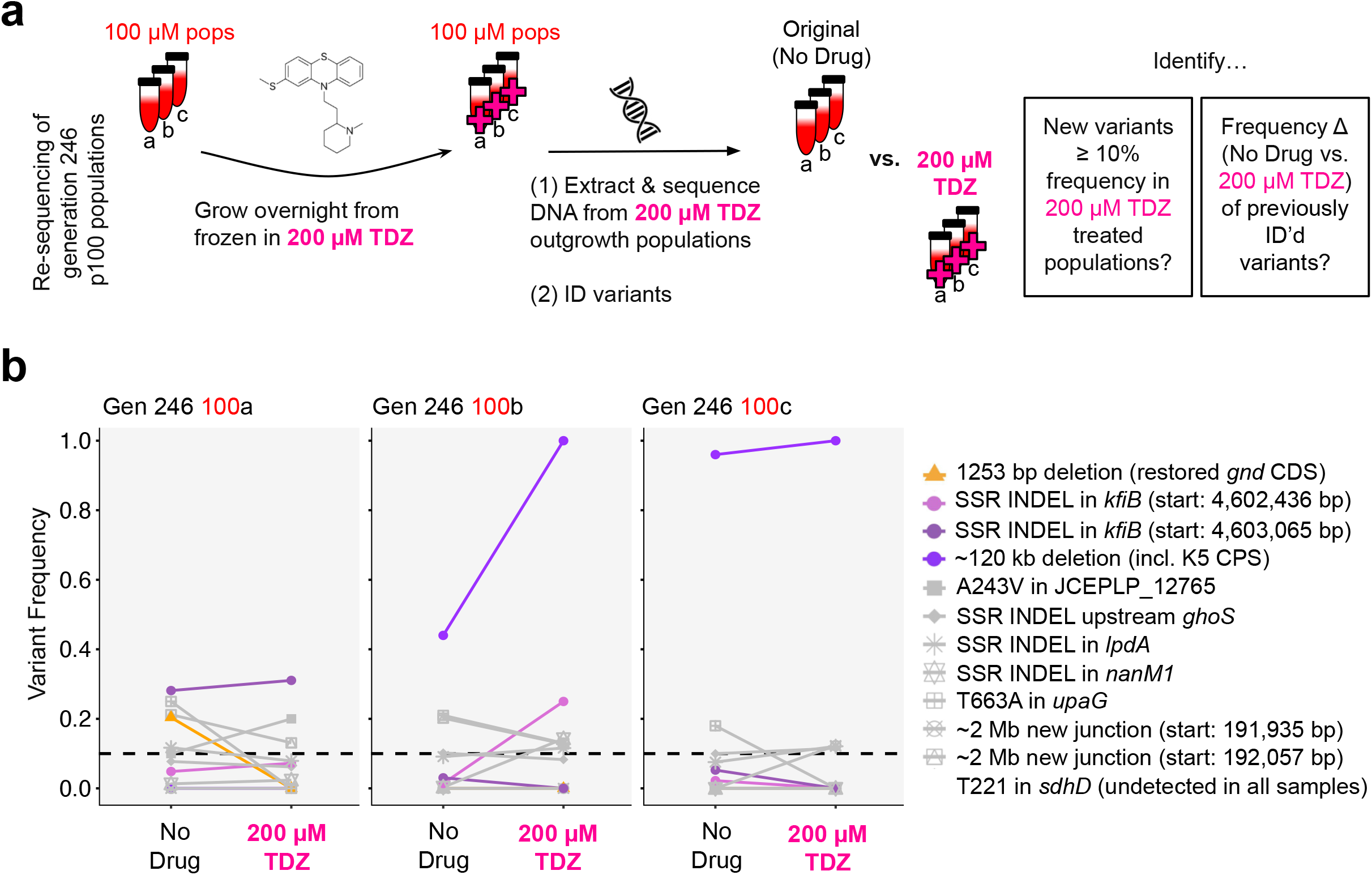
TDZ-resistant populations have elevated surface polysaccharide-linked variants under supra-MIC drug pressure. (**a**) Overview of the process for re-sequencing generation 246 p100 population replicates after reconstitution in M9-glucose spiked with 200 µM TDZ. Variants (from *breseq*, *PhaVa*, and the SSR-localized INDEL analysis) fitting at least one of two categories were assessed: those reaching ≥10% frequency in the 200 µM TDZ re-sequenced populations, and those that were already identified as variants of interest (Fig 2b). (**b**) Frequencies between the original (“no drug”) and 200 µM TDZ outgrowth population samples for the specified variants.

**Supplementary Figure 6.**
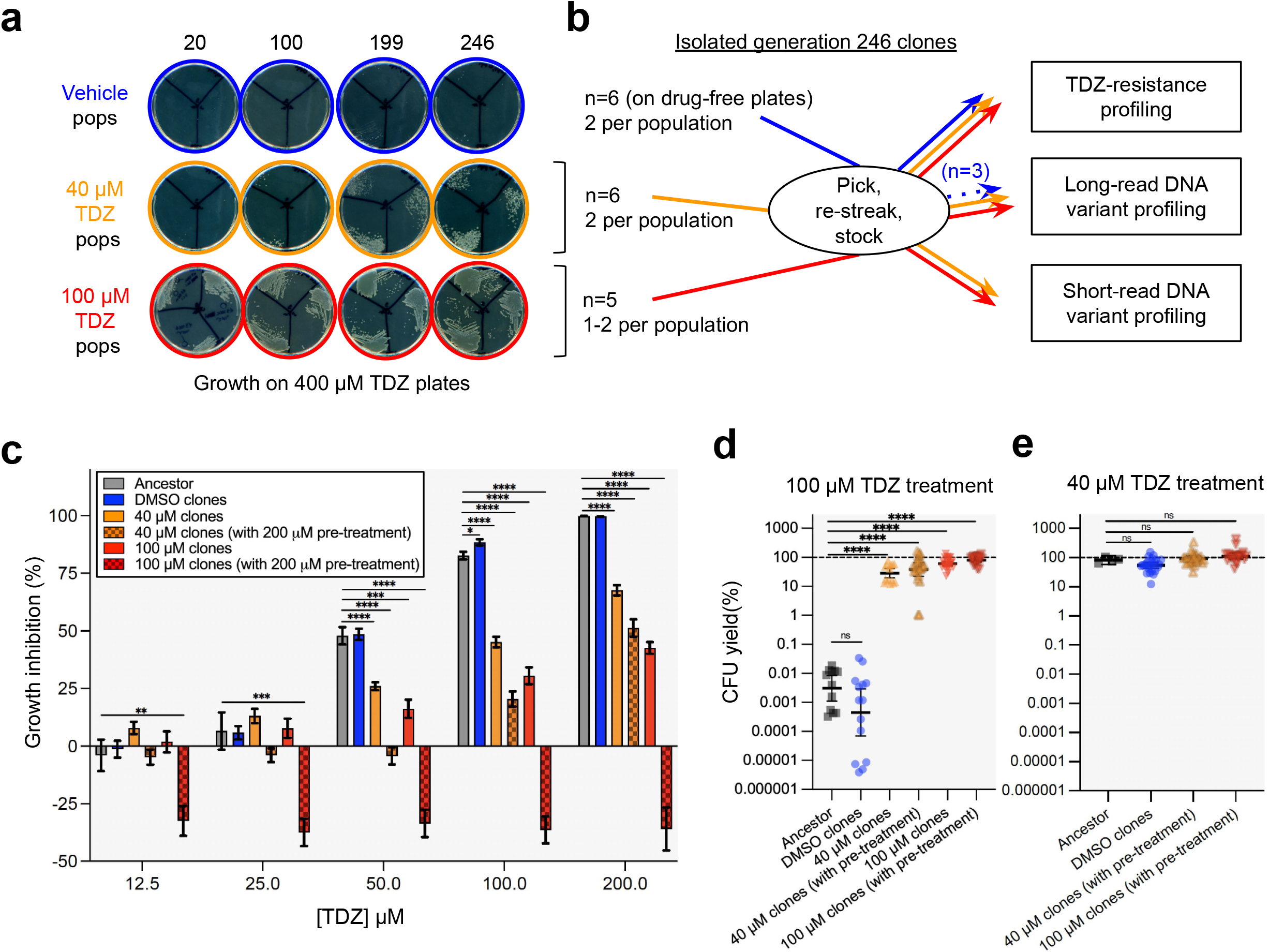
Both subinhibitory and colon-approximate TDZ exposure select for TDZ-resistant clones. (**a**) Contrast-enhanced images of glycerol stock-inoculated, 400 µM TDZ-spiked M9-glucose plates from each replicate population at generations 20, 100, 199, and 246. Replicate populations were plated in sections on the same plate. (**b**) Summary of the isolation and characterization process of generation 246-isolated clones. Both TDZ-sensitive (i.e. from pDMSO populations streaked on drug-free M9-glucose plates) and TDZ-resistant (i.e. from p40 and p100 populations streaked on 400 µM TDZ plates) were collected. (**c**) Dose-dependent growth inhibition (%) of the experimental evolution-derived clones compared to WT *EcN*, assessed by endpoint OD_600_ assays. TDZ-resistant clones were either grown with persistent TDZ exposure or in the absence of drug pressure immediately prior to growth inhibition assessment (see **Methods**). Data are mean ± SEM from four biological replicates for WT and two biological replicates for each of the six unique pDMSO clones, the six unique p40 clones (three without pre-treatment), and the five unique p100 clones (three without pre-treatment). All biological replicates were split into a minimum of two technical replicates. (**d**) CFU yield (%) after five hours of 100 µM TDZ treatment initiated at exponential phase. Error bars are the 95% CI centered on the geometric mean. Data are from the same number of unique clones and biological replicates as in **c,** except for ten pDMSO clone technical replicates falling below the limit of detection, and one of three unique p40 clones (without pre-treatment) which had colonies too small for CFU enumeration. (**e**) CFU yield (%) after five hours of 40 µM TDZ treatment initiated at exponential phase. Data are from the same number of unique clones and replicates as in **c**, but with only two biological replicates for WT and absent of p40-and p100-isolated clones without pre-treatment. Mann-Whitney U was used for comparisons: * = p < 0.05, ** = p < 0.01, *** = p < 0.001, **** = p < 0.0001.

**Supplementary Figure 7.**
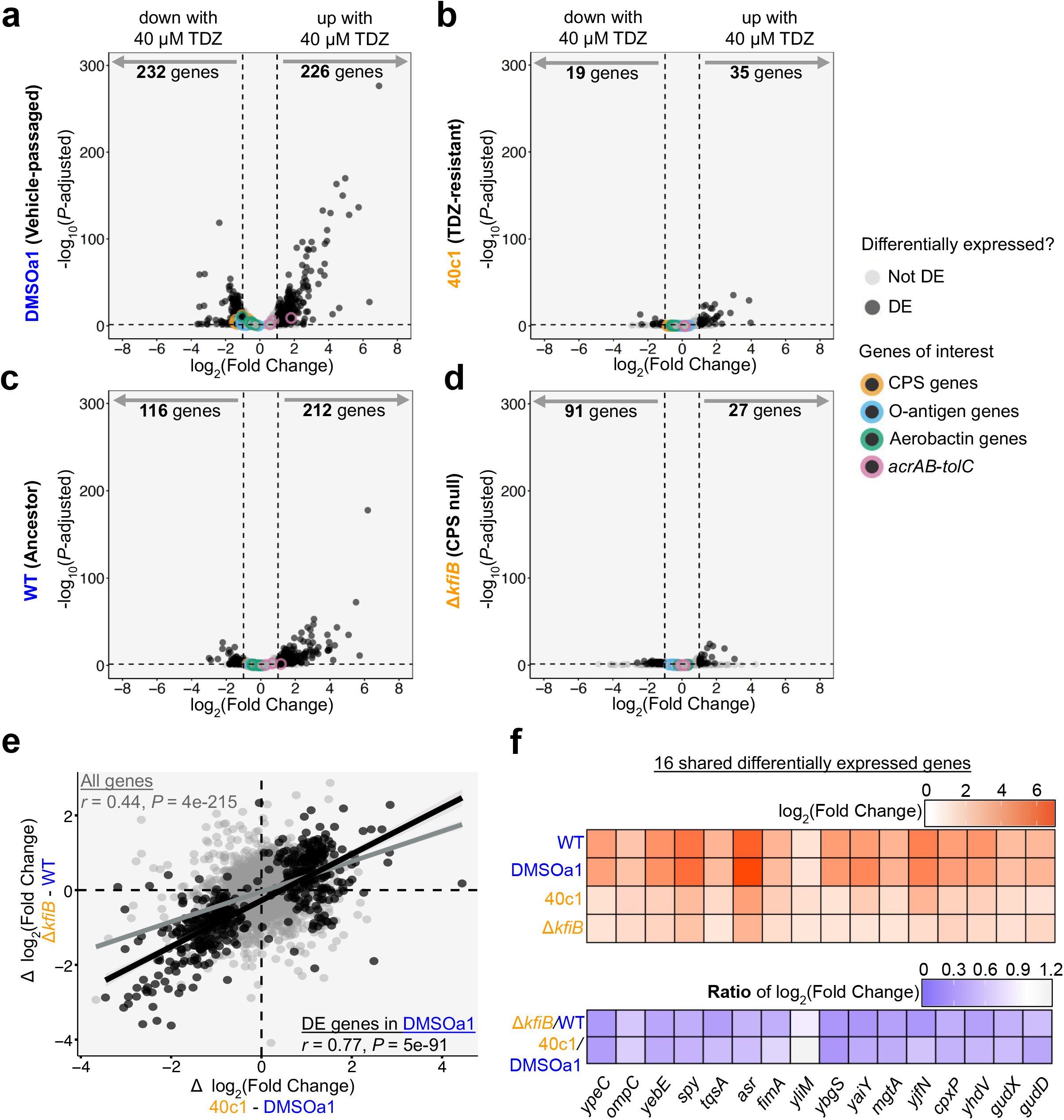
CPS expression sensitizes *EcN* to subinhibitory TDZ exposure. (**a-d**) Volcano plots of differentially expressed genes between vehicle (0.1% DMSO) and TDZ (40 µM) treatment for (**a**) DMSOa1, (**b**) 40c1, (**c**) WT, and (**d**) Δ*kfiB*. Differentially expressed (DE) genes fulfilled the following criteria: |log_2_(Fold Change)| ≥1.0 and *P-*adjusted <0.05, using the Benjamini-Hochberg correction for multiple hypothesis testing. (**e**) Scatterplot of the 4,665 genes (including non-rRNA ncRNAs) present in the 40 µM TDZ differential expression analysis across all four strains. The y-axis is the difference (Δ) in log_2_(Fold-Change) between the “unevolved” clones (WT subtracted from Δ*kfiB*), and the x-axis is the difference (Δ) in log_2_(Fold-Change) between the “evolved” clones (DMSOa1 subtracted from 40c1). Pearson correlation analysis was first performed across all genes (gray) and then across the 458 genes that were DE in DMSOa1 (black; only the 452 genes present in the differential expression analysis across all strains were included). (**f**) Log_2_(Fold Change) results for the 16 genes that were DE across all four strains (top) and their log_2_(Fold Change) ratios for Δ*kfiB*/WT and 40c1/DMSOa1 (bottom). The genes are in order of their position in the *EcN* reference genome.

